# Consistent covariation of dispersal, life history and thermal niche across terrestrial arthropods

**DOI:** 10.1101/2025.01.10.632369

**Authors:** Garben Logghe, Femke Batsleer, Guillermo Fandos, Dirk Maes, Dries Bonte

**Author notes:** **Corresponding author:** D. Bonte, Ghent University, Department of Biology, K. L. Ledeganckstraat 35, 9000 Ghent, Belgium. **Statement of Authorship**: Garben Logghe, Femke Batsleer, Dirk Maes and Dries Bonte conceived the framework of the study. Garben Logghe collected the trait data. Guillermo Fandos and Garben Logghe developed the statistical analysis. Garben Logghe wrote the first draft of the manuscript. All authors contributed substantially to interpretation of the results and revision of the manuscript. **Data accessibility statement**: The complete dataset on arthropod life histories and ecological traits can be found on GBIF: https://doi.org/10.15468/75g5z9. The filtered dataset specific to this study can be accessed on Zenodo: 10.5281/zenodo.14527806. Commented Rcode and dataset for brms-analyases on https://github.com/dbonte/TraitSyndromes_Logghe_etal.

## Abstract

Arthropods, as ectotherms, are experiencing global declines, with many species facing the need to either acclimate or disperse in response to climate change. Understanding to which degree life history, dispersal and thermal niche traits covary is key to improve distribution forecasting under climate change. We quantified life history, dispersal and thermal range covariation among 4000 Western European arthropod species spanning eight orders, considering phylogenetic relationships to account for common ancestry. We demonstrate the existence of two axes of life history variation: the fast-slow continuum and the reproductive strategy axis. Species at the fast end of the continuum have higher dispersal capacities and broader thermal niches than slower species. The resulting trait syndromes were surprisingly consistent across orders. These trait combinations, which generally enhance range-shifting potential, point to the emergence of two distinct groups of arthropods: those well-suited and those less equipped to mitigate the effects of future climate change.

## Introduction

Climate change is an escalating global problem, threatening numerous species across various taxonomic groups [1–3]. Species may cope with changing climatic conditions by acclimatisation and/or adaptation to the experienced novel conditions, or by tracking their climate window through dispersal [4]. The pace at which species expand or shift their distribution will critically depend on their life histories such as survival, reproduction and development, that determine population growth rate and overall demography. Furthermore, this rate of range shifts will be shaped by their dispersal capacity and ability to acclimate to novel conditions [4]. Forecasting biodiversity under climate change [5,6], therefore, relies on our ability to incorporate accurate trait data into extinction risk projections [7,8], thereby informing vulnerability assessments [9,10] and refining predictions of emerging community assemblages [11,12]. For arthropods, advancing trait-based forecasting is particularly urgent, as delays in benchmarking efforts risk exacerbating their steep global decline [13] and the associated loss of vital ecosystem services [14]. To date, most distribution models for arthropods focus on iconic or pest species and rely heavily on correlative approaches such as species distribution models (SDMs). Mechanistic models that are essential for predicting responses under novel environmental conditions remain rare, largely due to the lack of detailed data on species-specific life histories and physiological tolerances across these highly diverse taxa [15,16].

Species are expected to show intrinsic life history trade-offs, positioning them along a “fast-slow” continuum [17–19]. Slow-living species have longer development times and lifespans, prioritizing resource allocation toward offspring survival rather than fecundity, while fast-living species adopt the opposite strategy [20]. In theory, slow-living species are better adapted to stable environments and have, therefore, greater difficulty recovering from stochastic mortality events [21]. Due to similar evolutionary pressures stemming from environmental instability, dispersal rates are expected to be lower in species with slow life histories [22,23]. Body size, however, may have contrasting effects. While smaller body sizes are typically linked to a fast-paced lifestyle, larger body sizes are often positively correlated with greater dispersal ability [4,24–26]. Predicted relationships between species’ dispersal and life history traits are therefore highly variable, both among and within species [27].

Both dispersing towards new areas and staying within a changing environment comes with certain costs, such as the risk of staying or settling in environments where conditions may be unfavourable [28]. Such limitations should be especially pronounced in ecological specialists, as they will suffer most from local environmental changes and/or experience greater challenges in establishing populations in unfamiliar habitats [29–32]. In the context of climate change, thermal tolerance is a particularly important niche component in shaping distribution patterns [33]. A species’ thermal niche encompasses several critical factors, including its thermal minimum (the lowest temperature it can endure), thermal maximum (the highest temperature it can tolerate) and thermal range (the span between these extremes) [34]. Species with narrow thermal niche breadths—those with limited tolerance to temperature variation—are generally expected to be more vulnerable to climate change [35,36], facing greater risks during dispersal as they attempt to track their optimal temperature conditions. Recent findings from Neotropical rainforest species show that greater dispersal ability can reduce thermal specialization, thereby lowering extinction risk under climate change [33]. Given this pivotal role of a species’ thermal niche in determining both distribution ranges and dispersal costs, it can be expected to have shaped the evolution of both life histories [37,38] and dispersal [39,40].

Physiological tolerance to temperature, a key determinant of species’ climate sensitivity, can be inferred from occurrence data by linking georeferenced records to environmental variables such as temperature extremes. For taxa with good coverage and low misidentification rates, GBIF enables scalable, cost-effective assessments of thermal tolerance across space and time [41–44]. For example, historical species distribution models (SDMs) built from GBIF and museum data for Canadian butterflies predicted current ranges with high accuracy, demonstrating temporal niche stability [45]. Moreover, extracting environmental variables from GBIF records yields estimates consistent with IUCN expert maps, reinforcing its utility for macroecological inference [46].

Linking such occurrence-based insights with life history strategies provides a more holistic framework for predicting species’ responses to climate change. General trait syndromes (i.e. combinations of traits that covary predictably across taxa) can reveal how life history, dispersal and thermal niche interact to shape species’ resilience. Traits underlying life history variation can show complex covariation [47–49], and dispersal syndromes, which document covariations between dispersal and other traits, have been documented across various taxonomic groups [22,23,50–52]. Such trait relationships may have important implications for species’ vulnerability to climate warming. For example, a positive correlation between dispersal ability and thermal niche breadth would suggest interspecific differences in vulnerability, as species with high dispersal capacity and broad thermal niche are likely to be more resilient [33]. In contrast, a negative correlation between these traits could indicate trade-offs that limit persistence–enabling species to respond through either range shifts or thermal tolerance, but not both. Similar advantages or trade-offs may also emerge from covariation of other traits.

Understanding the organisation of trait syndromes not only allows us to estimate missing trait data in understudied taxa [23,53], but also allows more accurate forecasting of species responses to environmental change [54]. Urban et al. [55] argue that missing trait values need not preclude forecasting; instead, traits can be inferred from accessible data, enabling broader and more efficient predictions. When trait syndromes are known, virtual species approaches with demography and dispersal linked to well-documented master traits like body size, become powerful tools. This has, for instance, been demonstrated for terrestrial mammals in previous work that simulated realistic organisms with evolved trait combinations [56]. This shift allows theory to better reflect ecological realism compared to broad theoretical approaches (e.g., [57]), enhancing the utility of simulations for conservation and policy planning.

This study aims to quantify covariation between dispersal, life history, and thermal niche traits across Western European arthropods using phylogenetically corrected regressions. We hypothesize that (1) trait relationships vary across taxa due to morphological and ecological diversity, and (2) dispersal and thermal range traits align with the fast-slow life history continuum, enhancing its applicability in global change forecasting.

## Material and Methods

### Data description

The data used for this study was sourced from the dataset described by Logghe et al. (2025), which contains information on 28 life history and ecological traits for 4874 terrestrial arthropod species. We focused on eight arthropod orders for which data on both dispersal and life history was available from a total of 83 different literature sources, complemented by unpublished measurements from taxonomic experts: Araneae, Coleoptera, Hemiptera, Hymenoptera, Isopoda, Lepidoptera, Odonata and Orthoptera. Two orders, Diptera and Opiliones, were excluded because of data shortage and the predominant reliance on expert judgment. This resulted in a dataset containing 4343 species. From this dataset, we selected nine traits related to demography (life history), dispersal (movement) and physiology (thermal niche), relevant for predicting range shifts under climate change [55,59]. Appendix S.1 shows the number of data points for every trait within each order.

We selected body size as a central trait, since it is shown to correlate with life history, dispersal ability and thermal limits [60–62]. In this study, we use the adult body length (measured in millimetres) as a proxy for body size. In contrast to several other highly correlated, relevant measures of body size (i.e., body mass, volume; [63,64], body length is always reported in the primary literature. To avoid introducing additional and unquantifiable uncertainty in our analyses, we retained this metric rather than transforming it to one of the derived metrics that require assumptions about body shape that are not consistently documented, even within taxonomic orders.

As demographic traits, we selected development time, fecundity and voltinism. Development time is defined as the duration between egg laying and adult emergence, expressed in days. Fecundity is defined as the number of offspring a female could produce during her lifetime, while voltinism refers to the maximum number of generations a species could complete in a single year. Note that the maximum value for voltinism is used, as the number of generations can vary annually and across regions. All these trait values were obtained from primary literature (see Logghe *et al*. [58]).

We collected mobility-related trait data that serve as well-known proxies for dispersal capacity (the ability to disperse – wing load, wing morphology in most flying orders), dispersal motivation (ballooning propensity in spiders), movement speed (Isopoda) and mean dispersal distances (various groups) as reported from literature and unpublished data from experts in Logghe et al. [58]. To enable cross-species comparison, we standardized dispersal scores on a scale from 0.1 (highly philopatric) to 1 (highly mobile). Importantly, these scores are relative within each order, meaning that a butterfly and a spider with a score of 1 may differ substantially in absolute dispersal capacity, but both represent the most dispersive species within their respective groups. This transformation translates the different proxies into a single measure for dispersal motivation - the propensity to disperse (or remain philopatric). Since dispersal distance kernels are of the inverse power-law family, high values of our proxy would also translate into a higher frequency of individuals engaging in further dispersal distances, supporting its ecological relevance [65,66].

Finally, we included measures of each species’ thermal niche, specifically thermal mean and range (the difference between the thermal minimum and maximum). These metrics were derived by overlaying GBIF species distribution data with a raster of climate records, allowing us to assess the climate conditions under which species currently occur. Using zonal statistics for raster values within defined geographic zones, we calculated the average, lowest and highest annual temperatures (in °C) experienced across each species’ geographic range (see Logghe *et al*. [58] for technical details). As distribution data were obtained directly from GBIF, our estimates represent the realised thermal niche, defined here as the range of temperature conditions a species experiences within its known distribution. While using GBIF data may introduce observer bias, we expect this bias to be reduced in our study, as we focus on well-documented species that are more likely to have reliable and comprehensive trait data available. This approach does not necessarily capture the full physiological (or fundamental) thermal limits of a species. Instead, it reflects the environmental temperatures species currently encounter, so the conditions that may be shaped by factors such as dispersal constraints or biotic interactions. However, we intentionally excluded species that are endemic to certain geographically distinct locations. Ranges of species strictly engaged in host-specific interactions may be primarily constrained by the distribution of the host rather than by its own thermal niche. We did not include such highly specialised species, and the few considered are documented to shift hosts within their geographic range (e.g. more specialist herbivores or parasitoids). Although these trait data are assumed to strongly link to thermal optima and niche, we further refer to them as thermal mean and thermal range.

It is important to note that this study does not account for intraspecific variation in trait values. The literature mostly reports mean values, occasionally accompanied by minimum and maximum ranges, but rarely with proper variability statistics. We thus developed a species-level meta-analysis, using mean trait values to explore broad patterns of trait covariation across the phylogenetic tree.

### Statistical analysis

R version 4.0.4 [67] was used to perform all data analyses. We used tidyverse [68] for data manipulation and plots were made using ggplot2 [69].

#### Constructing the phylogenetic tree

To conduct phylogenetic analyses and apply corrections, we required a phylogenetic tree encompassing all species in the dataset to serve as foundation for subsequent analyses. The tree was built by linking species names to the Open Tree Taxonomy database, which contains data on known phylogenetic relationships (“rotl” package) [70]. Branch lengths were added to the tree using the “ape” package [71] and calculated with Grafen’s method [72], which transforms a cladogram into an ultrametric tree by scaling internal node heights according to the number of descendant taxa. This approach assumes a speciational model of divergence, where evolutionary change is linked to speciation events rather than elapsed time. The resulting tree is appropriate for use with comparative methods that assume Brownian motion trait evolution (i.e., traits evolve via a random walk with constant variance over time).

#### Phylogenetic signals

Shared evolutionary history may result in closely related species showing similar trait combinations due to their shared ancestry and not because of the ecological link between the traits [73]. Such similarities between related species can potentially obscure or exaggerate recurring patterns, reflecting shared ancestry rather than independent trends. We assessed the phylogenetic signal using Pagel’s Lambda for each considered trait and arthropod order [74]. This is a metric that estimates how much closely related species resemble each other compared to expectations under a Brownian motion model of evolution [75]. Pagel’s Lambda typically ranges from 0 (indicating no influence of phylogenetic relationships on trait values) to 1 (indicating trait values are entirely explained by phylogenetic relationships). We calculated this metric using the “phylosig” function from the “phytools” package [76]. We visualised the distribution of each assessed trait across circular phylogenetic trees of terrestrial arthropods and represent size, thermal range, fecundity and body size in the main manuscript. To facilitate comparison between the trees, we selected only species with data available for all four traits (n = 297). Fecundity and size were log-transformed to improve clarity of the figure by creating a smoother colour gradient.

#### Identifying trait syndromes

To account for the influence of shared evolutionary history on trait similarities, we controlled for phylogenetic relatedness in all analyses. Such corrections were essential to meet the assumption of statistical independence, as closely related species often show greater similarity due to shared ancestry. This approach also helps confirm that the detected patterns occur within clades, not just between them. For example, suppose trait A shows large variation between clades (e.g. high in one clade, low in another), and trait B varies similarly within each clade. In this case, an apparent correlation between traits A and B may emerge, primarily driven by between-clade differences. Phylogenetic models help separate the within- and between-clade variation, revealing whether trait correlations reflect repeated patterns within lineages or are merely the result of shared ancestry. Since this phylogenetic correction is subject to assumptions on ancestry [73], we also provide uncorrected estimates in Appendix S.3 to assess how much the retrieved correlations are sensitive to the implemented model of evolution.

Bayesian Phylogenetic Generalized Linear models were employed to assess whether covariations were present between the selected traits (“brms” package) [77]. Given the strong phylogenetic signal across the tree of life, the variation in data quantity and quality among orders, and our aim to detect and understand potential divergence in the covariations across arthropod orders, we analysed every order separately. Trait data on body size, fecundity and development time underwent log transformation before analysis to reduce the skewness of the data. Because we had no a-priori causal mechanism in mind, each trait combination was run twice, with the dependent and independent variables interchanged. This dual approach revealed consistent statistical significance of the pattern (see Appendix S.4). Given the strong phylogenetic signals among orders, we chose not to construct a single multivariate model, also to avoid potential biases arising from uneven patterns of missing data among taxa. Additionally, separate models allowed us to account for different error structures, depending on the nature of each trait. Each model was, therefore, fitted with a distribution appropriate to its independent variable: zero-one-inflated beta for voltinism, binomial for dispersal and Gaussian for the remaining variables. The models were run using 4 chains and 4000 iterations, with a burn-in period of 1000. Non-informative flat priors were used for the model parameters. Phylogenetic relationships between species were incorporated as a species-level effect with a covariance matrix derived from the previously mentioned phylogenetic tree (“geiger” package) [78]. Specifically, we included the term (1|gr(species, cov=A)), where A is the phylogenetic covariance matrix assuming Brownian motion evolution. Convergence of the models was checked by examining the effective sample size (>1000) for each chain as well as Gelman-Rubin statistic (Rhat < 1.1) among chains. Effect sizes were deemed clearly determined when 0 was not included in the 95% credibility intervals of the posteriors.

To examine how life history, dispersal ability and thermal traits are distributed across arthropod orders, we performed a phylogenetic principal component analysis (PCA) using the “phyl.pca” function from the “phytools” package [76]. Unlike standard PCA, this approach incorporates the phylogenetic covariance matrix, which reflects the expected covariance of traits under a Brownian motion model of evolution. The analysis was conducted on a subset of the dataset that included only the 200 species for which complete data on all traits were available. Note that data standardisation does not affect the ordination of the species according to their traits, and that this phylogenetic correction allows us to detect consistencies among trait correlation across taxa.

## Results

### Phylogenetic signals

We provide an overview of the phylogenetic signals (Pagel’s Lambda) per taxonomic order and trait in Appendix S2, which are summarised in Figure 1.

**Figure 1.**
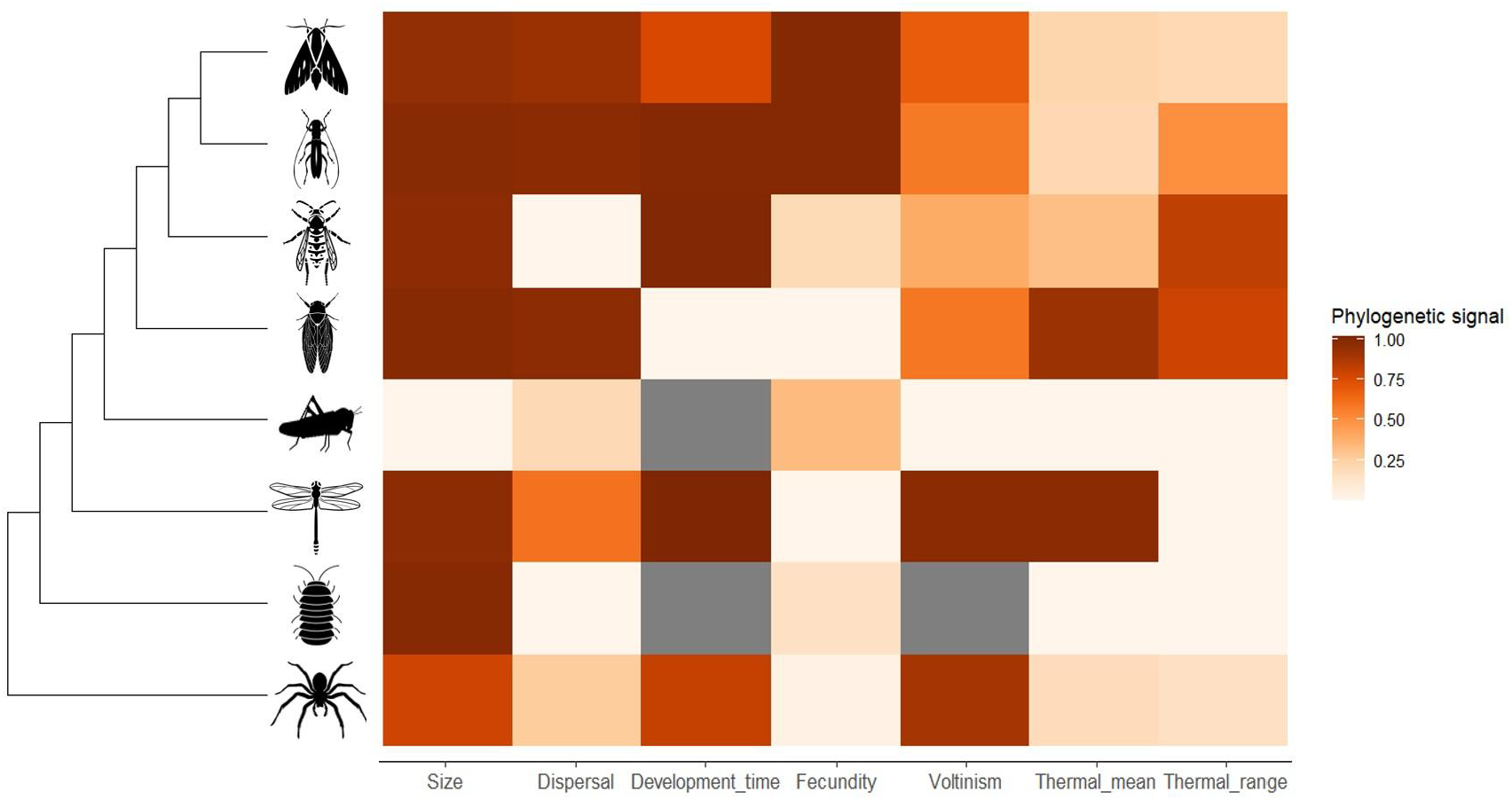
Overview of the phylogenetic signal (Pagel’s λ) for every assessed trait for each arthropod order. Arthropod orders are from top to bottom: Lepidoptera, Coleoptera, Hymenoptera, Hemiptera, Orthoptera, Odonata, Isopoda and Araneae. Colours represent the strength of phylogenetic signal, with higher values (darker red) indicating that trait values are more explained by phylogenetic relationships. The grey squares indicate that the phylogenetic signal was not calculated for that trait and order due to a lack of data. Sample sizes for every order and trait can be found in Appendix S.1.

The distributions of body size, dispersal, fecundity and thermal range are given in Figure 2. Across taxonomic orders, body size consistently shows a strong phylogenetic signal (Pagel’s λ > 0.78), while traits like fecundity and thermal range exhibit more variable or weak signals. Orders such as Coleoptera and Lepidoptera display high trait conservatism across most traits, whereas Orthoptera and Isopoda show minimal phylogenetic structuring for thermal and dispersal traits. The phylogenetic signal for dispersal displayed considerable variation, indicating that the influence of phylogenetic relationships differs substantially between orders.

**Figure 2.**
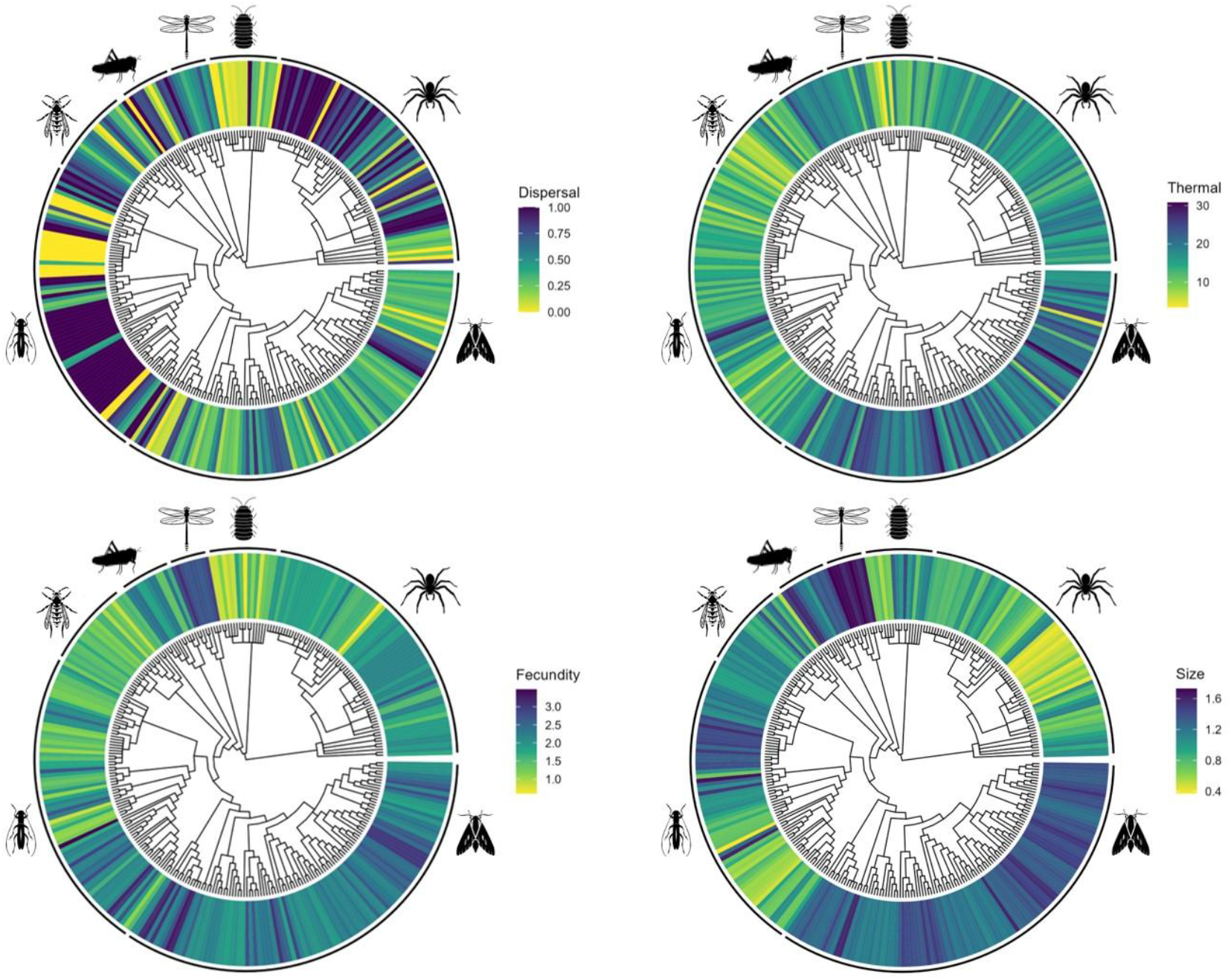
Overview of the distribution of trait values across the phylogenetic tree for dispersal (λ = 0.983), fecundity (λ = 0.988), thermal range (λ = 0.863) and body size (λ = 0.993). To facilitate comparison between the trees, we selected only species with data available for all four traits (n = 297). Fecundity and Size were log-transformed to improve clarity of the figure by creating a smoother colour gradient.

### Identifying trait syndromes

Our results reveal multiple trait syndromes across various arthropod orders (Figure 3, Appendix S.4). In most cases, voltinism is positively correlated with dispersal capacity. Furthermore, temperature niche components generally covary positively with dispersal capacity, fecundity and body size across most orders. Body size typically shows a negative relationship with voltinism, and a positive one with dispersal capacity, although Coleoptera represent a notable exception. Overall, size, dispersal, fecundity and thermal traits show consistent patterns of covariation across orders. While other trait associations also emerge within individual orders, they appear to be less broadly generalizable. However, it should be noted that, aside from a few exceptions, most orders show either similar correlation directions to the others or no clearly determined relationship at all.

**Figure 3.**
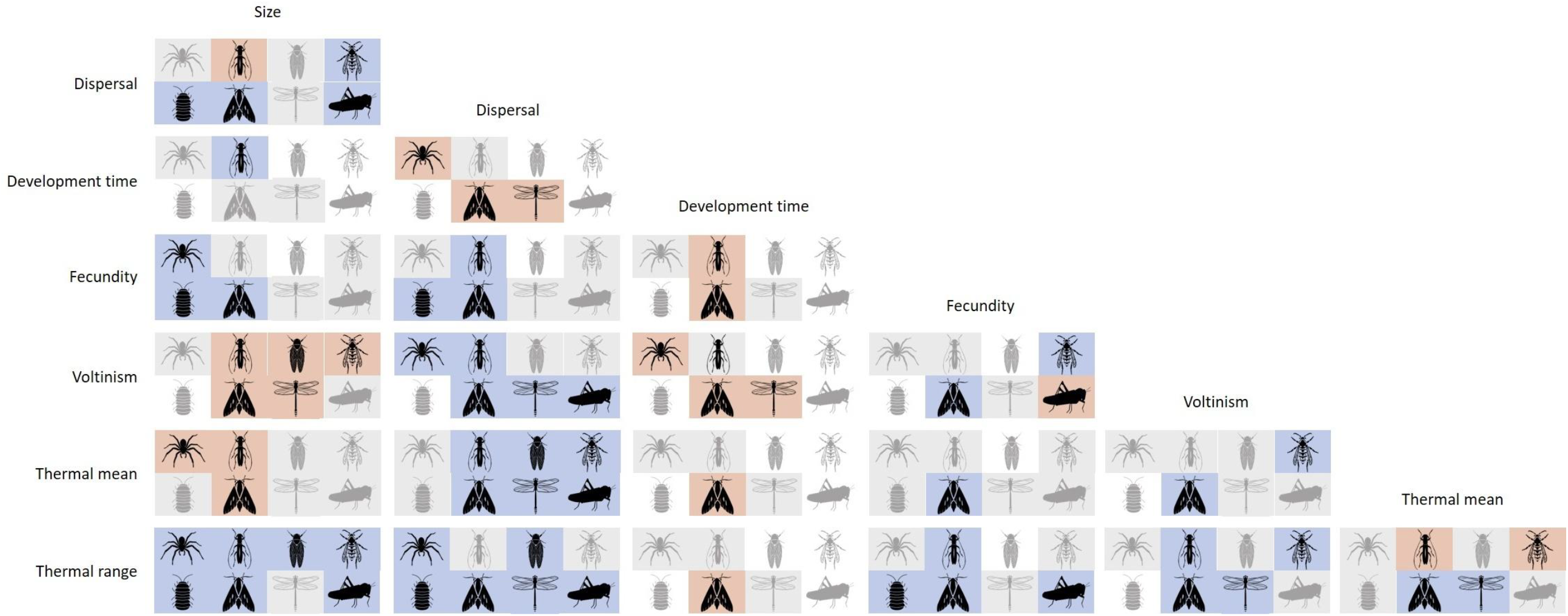
Overview of trait covariations across each order, represented as a correlation matrix. Each grid of the matrix is divided into eight cells, with each cell representing one order. The taxa are, from left to right and top to bottom: Araneae, Coleoptera, Hemiptera, Hymenoptera, Isopoda, Lepidoptera, Odonata and Orthoptera. Red cells indicate a clearly determined (0 not included in the 95 credibility interval) negative relationship, blue cells indicate a clearly determined positive relationship, and grey cells show that the relationship was not clearly determined. White cells denote that this trait combination was not assessed for the given order, due to either an absence of data for one of the traits or convergence issues. Sample sizes for every covariation were determined by the trait with the least available data (see Appendix S.1).

The pairwise models incorporated phylogenetic relationships between species, which helps avoid biased results caused by treating closely related species as independent. Although the traits showed strong phylogenetic signals (Figures 1–2), the direction of trait correlations remained consistent across models that accounted for evolutionary history using a Brownian motion model and the available phylogenetic tree, and those that did not (see Appendix S.3). However, in many cases, the credibility intervals of the phylogenetically corrected models included zero, indicating weaker statistical support for these correlations compared to models that did not correct for phylogeny (Appendix S.3).

The phylogenetically corrected PCA revealed no distinct separation among orders, confirming that covariations are overall consistent across taxonomic groups (Figure 4). PC1 shows a strong positive relationship with voltinism and dispersal, while it also strongly negatively correlates with development time (Appendix S.5). Additionally, it exhibits a positive correlation with thermal mean and range, though this relationship is not as strong as with the other traits. In contrast, PC2 is strongly positively correlated with fecundity and size. These findings support the previous results that correlations between body size, dispersal, thermal traits and voltinism observed within individual orders hold across multiple arthropod orders.

**Figure 4.**
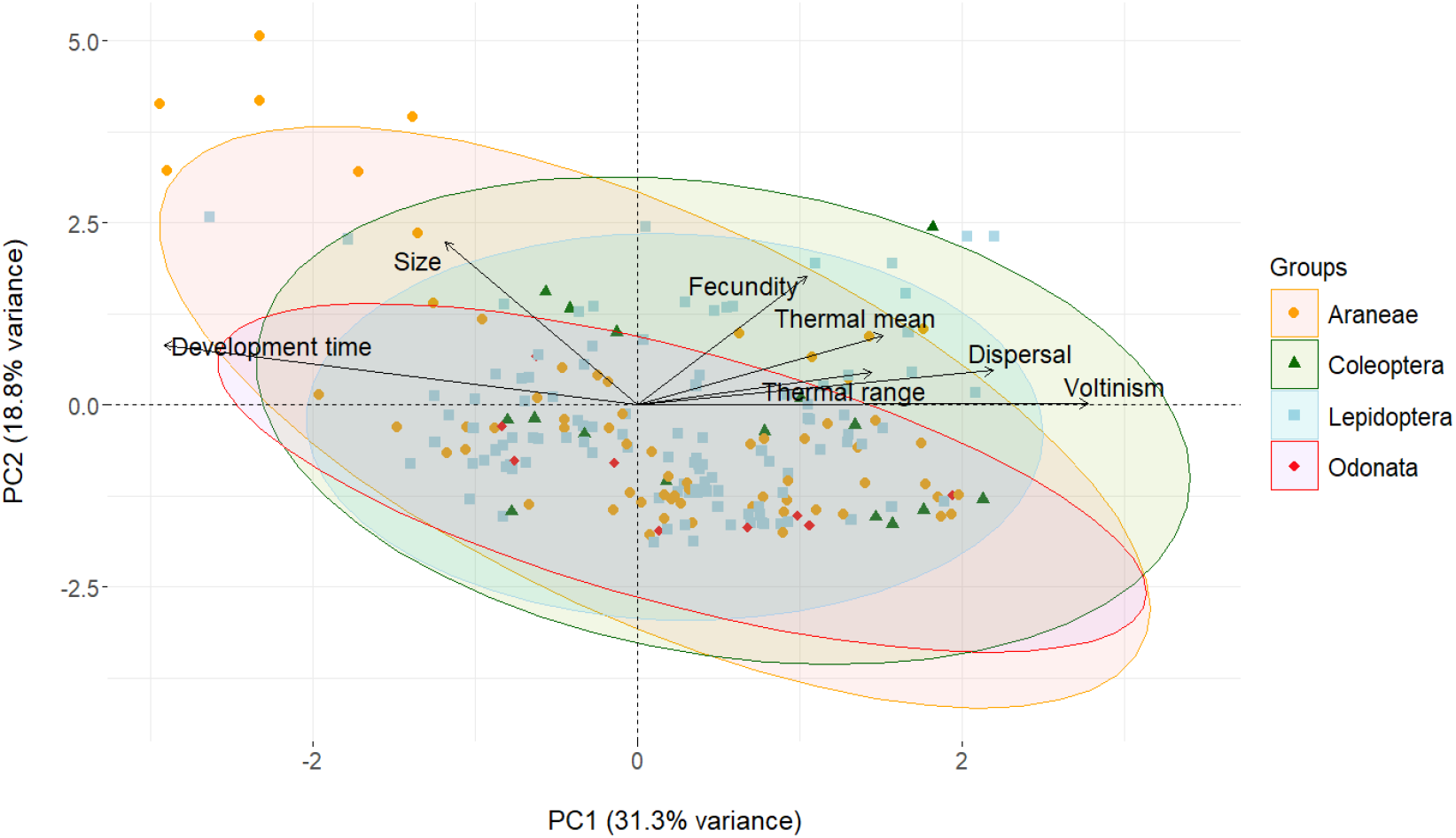
Phylogenetically corrected PCA results for arthropod species from the four most important orders for which data on all traits was available (n=200). Dots represent species with symbol colour and shape according to their taxonomic order. Arrows represent the direction and magnitude of each variable’s contribution to the principal components. Ellipses represent 95% confidence intervals around species scores for each arthropod order, illustrating group-level patterns in trait space.

## Discussion

Our quantitative synthesis across a wide range of Western European arthropods indicates that dispersal, life history and thermal niche (mean and range) are tightly connected, implying similar evolutionary pressures on these traits. Although phylogenetic signals were quite variable across orders and traits, some traits, such as body size and development time, show a strong influence of shared ancestry on trait expression. This confirms the importance of phylogenetically corrected analyses for identifying trait syndromes. While there are some notable differences in the trait syndromes between orders, these relationships remain largely consistent across taxa, which is contrary to our expectation. In line with our second expectation, dispersal and thermal range show to be tightly linked with life history traits and integrated within the fast-slow continuum.

The relatively large consistency of trait syndromes across orders suggests that trait covariations can be generalized to a broad range of arthropod species. This remarkable consistency across arthropod orders is one of the most surprising findings of this study, given the extraordinary diversity of dispersal mechanisms and life history strategies within the phylum. Such uniformity could have important implications for modelling the future effects of climate change, suggesting that detailed trait information for every species might not be as necessary as previously suggested [55]. Instead, missing trait values could potentially be inferred from accessible trait data, facilitating broader, more efficient predictions. For mechanistic modelling, traits evidently need to be thoughtfully translated to demography and the emerging population dynamics or ecosystem properties [16].

A considerable amount of interspecific life history trait variation in arthropods can be attributed to the fast-slow continuum, with a clear distinction between species with fast (i.e., short development times) and slow (i.e., long development times) lifecycles. Interestingly, development time appears closely tied to dispersal capacity, as we observed clear covariations between development time, voltinism and dispersal across various orders. Previous research has shown similar patterns, with high dispersal ability often linked with fast life history strategies [22,23,79]. Given that most insects disperse only in their winged adult stages, this dispersal-life history syndrome may have evolved as a bet-hedging strategy in response to unstable environments, allowing for rapid relocation when conditions become unfavourable [80,81]. Interestingly, we found limited evidence to suggest that fast-living species produce more offspring, a pattern that would typically be anticipated within the fast-slow continuum framework [20]. While Lepidoptera and Coleoptera did show a clear negative relationship between fecundity and development time (Figure 3), our analyses suggest that the reproductive axis of variation is as good as independent from the developmental axis. We thus show that the independence of these two distinct axes of life history variation is rather a rule than an exception, as was previously reported for plants [82] and vertebrates [18,83,84].

Our results suggest that thermal range has played a role in shaping the evolution of dispersal in arthropods, with most orders showing a positive relationship between thermal range and dispersal ability. This relationship likely arises from the costs associated with dispersing to new environments, where the risk of encountering unfavourable climatic conditions is high [28]. A broader thermal range reduces this risk by increasing the chances that newly colonised areas fall within the species’ tolerable thermal range. However, the causal relationship could also work in reverse, as limited dispersal capacity may constrain a species’ realised thermal niche by restricting access to potentially suitable regions, despite physiological tolerance to those climates [85]. In other words, the realised thermal range, as considered in this study, would be narrower than the fundamental thermal niche width, simply because of dispersal limitations. Implementation of other niche components such as diet or habitat specialisation could have provided a more holistic view on the relationship between dispersal and ecological generalisation. It is worthwhile to notice that the retrieved thermal range-body size relationships were predominantly negative, which aligns with experimentally derived patterns of lower thermal tolerance in large-bodied animals [86].

Besides thermal niche range, thermal mean values also covary with dispersal ability. Across nearly all assessed arthropod orders, species with distributions in warmer climates exhibit higher dispersal capacities. Although cold-adapted species may face greater risks from climate change and thus might benefit more from range shifting as a mitigation strategy, their generally lower dispersal capacities make spreading a less viable option [87]. Furthermore, our findings indicate that low dispersal capacity often covaries with slower life histories, which hampers colonization success due to slower population growth rates and, therefore, elevated vulnerability to stochastic extinction [88]. Consequently, the arthropod species most at risk from temperature changes due to climate change are also those least equipped to mitigate its effects through range shifts, a trend also noted in European mammals [89]. Overall, these consistencies of thermal niche correlations show that the proxies as derived from GBIF data are not randomly distributed and integrated into life histories according to predictions from the first principles of thermal ecology.

Phylogenetic signals in arthropod traits varied widely across orders and traits, with body size and development time showing strong conservation, as expected due to their ontogenetic linkage [90,91]. Interestingly, only Coleoptera showed a clear positive relationship between size and development time, suggesting that pace-of-life strategies may be more tightly conserved than direct trait correlations. In contrast, thermal niche traits exhibited weak phylogenetic signals, consistent with strong directional selection driven by environmental conditions rather than shared ancestry [92–95]. Dispersal and fecundity traits showed mixed patterns: some orders, like Coleoptera and Hemiptera, displayed strong phylogenetic structuring due to traits like wing development [96–98] and or a general body-size dependency [26,99,100], while others, such as Araneae, showed low phylogenetic influence, with dispersal more closely tied to ecological factors like habitat stability and rarity [101]. Overall, the observed phylogenetic signals align well with theoretical expectations, likely reflecting both evolutionary constraints and ecological adaptations.

Despite the variable phylogenetic signals of the different trait values, trait covariation is surprisingly consistent, even after controlling for signals of shared ancestry. This implies that relationships among traits are not only in the same direction across related species, but that they hold across less-related species from the same order, and even among orders. Our analyses support the existence of generalisable trait syndromes, suggesting that key ecological traits such as dispersal capacity or thermal range can be inferred from more readily available traits like body size. This consistency provides a valuable framework for predicting species persistence under environmental change, particularly for understudied taxa, and reinforces the utility of trait-based approaches in biodiversity forecasting. However, it is important to note that these findings are based on Western European species, where trait and distribution data are relatively well-documented. In more species-rich and data-sparse regions, such as the tropics, demographic and physiological traits remain poorly recorded. To extend trait-based forecasting globally, greater efforts are needed to collect trait proxies and occurrence data for hyper-diverse taxa and species diversity hotspots, ensuring that biodiversity models are inclusive and ecologically realistic.

## Acknowledgements

The authors would like to thank all contributors to the original dataset from which this study’s data were sampled: Tristan Permentier, Matty P. Berg, Pallieter De Smedt, Jonas Hagge, Jorg Lambrechts, Marc Pollet and Fons Verheyde. The computational resources (Stevin Supercomputer Infrastructure) and services used in this work were provided by the VSC (Flemish Supercomputer Center), funded by Ghent University, FWO and the Flemish Government – department EWI. Finally, the authors extend their gratitude to the Flemish Research Fund for their support of the Scientific Research Network EVENET (W001322N). All arthropod icons used in this paper originate from Noun Project (CC BY-NC-ND 2.0): Spider by Matthew Davis, Beetle by Rachel Siao, Cicada by Alejandro Capellan, wasp by Parkji Sun, isopod by Pham Thanh Lôc, Moth by Parkji Sun, Dragonfly by Hermine Blanquart and Cricket by Ed Harrison.

## Ethics declarations

The authors declare that they have no competing interests.

