## Supplementary material for "Consistent covariation of dispersal, life history and thermal niche across terrestrial arthropods"

#### S.1 Data coverage of traits

*Table S.1: Overview of the number of data points per trait and order. Data on body size was available for every species, meaning that this value also represents the total number of assessed species within each order.*

| Order | Size | Dispersal | Fecundity | Development time | Voltinism | Thermal |
|---|---|---|---|---|---|---|
| Araneae | 439 | 414 | 72 | 83 | 155 | 426 |
| Coleoptera | 1370 | 1321 | 87 | 238 | 347 | 1342 |
| Hemiptera | 708 | 628 | 26 | 27 | 508 | 667 |
| Hymenoptera | 567 | 86 | 53 | 38 | 399 | 523 |
| Isopoda | 34 | 29 | 10 | 0 | 5 | 34 |
| Lepidoptera | 1110 | 173 | 129 | 111 | 917 | 1054 |
| Odonata | 69 | 63 | 12 | 52 | 63 | 68 |
| Orthoptera | 51 | 46 | 7 | 0 | 33 | 50 |

### S.2 Phylogenetic signals

*Table S.2: Overview of the phylogenetic signals (Pagel's Lambda) per taxonomic order and trait.*

| Order | Size | Dispersal | Fecundity | Development time | Voltinism | Thermal mean | Thermal range |
|---|---|---|---|---|---|---|---|
| Araneae | 0.786 | 0.264 | 0.035 | 0.809 | 0.890 | 0.182 | 0.155 |
| Coleoptera | 0.982 | 0.978 | 0.997 | 0.996 | 0.580 | 0.209 | 0.498 |
| Hemiptera | 0.989 | 0.973 | < 0.001 | < 0.001 | 0.589 | 0.922 | 0.792 |
| Hymenoptera | 0.967 | < 0.001 | 0.193 | 1.001 | 0.388 | 0.319 | 0.820 |
| Isopoda | 0.986 | < 0.001 | 0.155 | N/A | N/A | < 0.001 | < 0.001 |
| Lepidoptera | 0.946 | 0.927 | 0.996 | 0.762 | 0.680 | 0.220 | 0.204 |
| Odonata | 0.970 | 0.603 | < 0.001 | 1.013 | 0.976 | < 0.001 | < 0.001 |
| Orthoptera | < 0.001 | 0.200 | 0.332 | N/A | < 0.001 | < 0.001 | < 0.001 |

### S.3 Comparison between corrected and uncorrected models

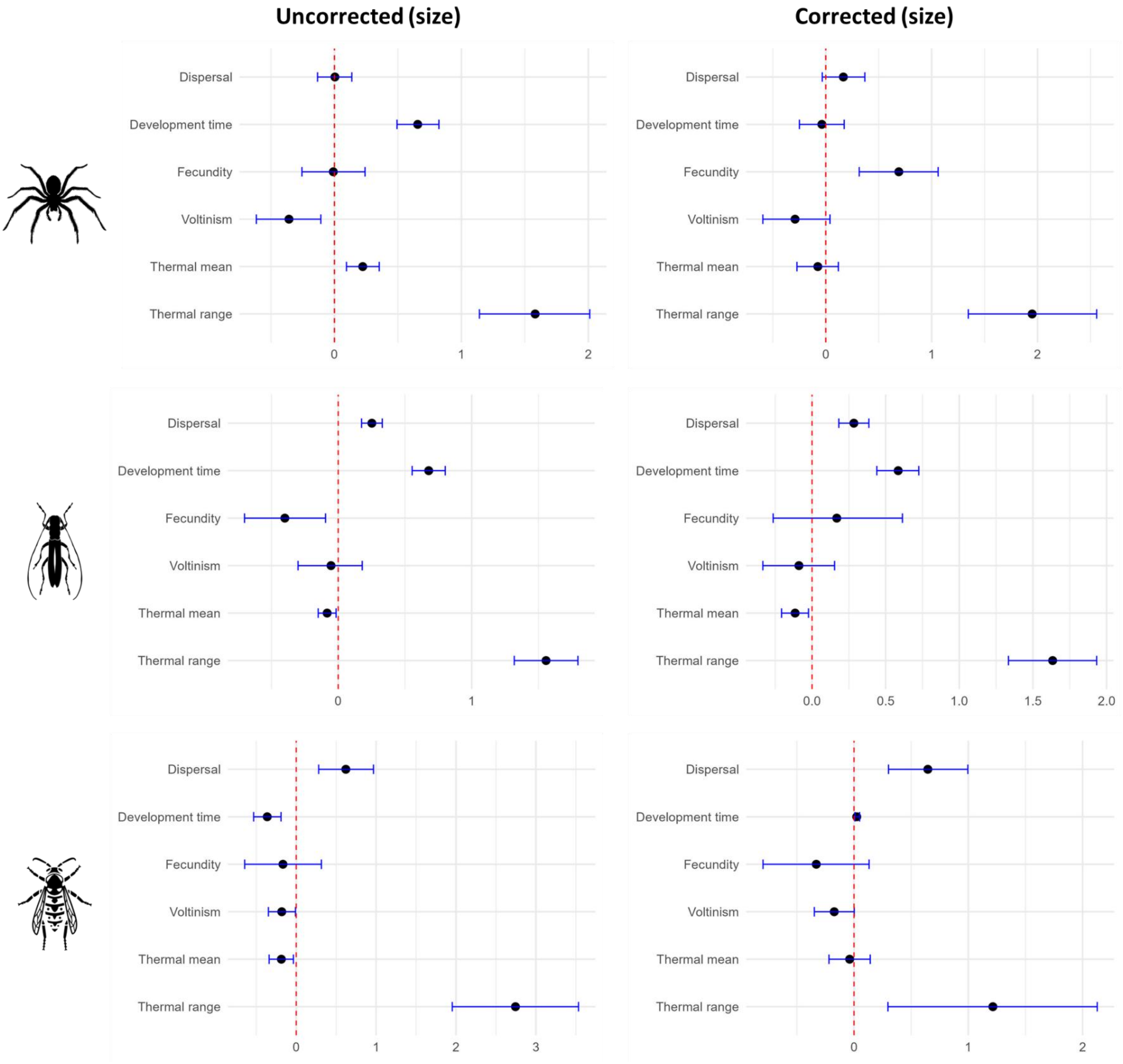

*Figure S.3.1a: comparison of uncorrected (left) and phylogenetically corrected (right) correlations with size for spiders, beetles and hymenopterans. Each point represents the correlation estimate with size as the independent variable and the corresponding trait as dependent variable. The error bars represent the 95% credibility intervals.*

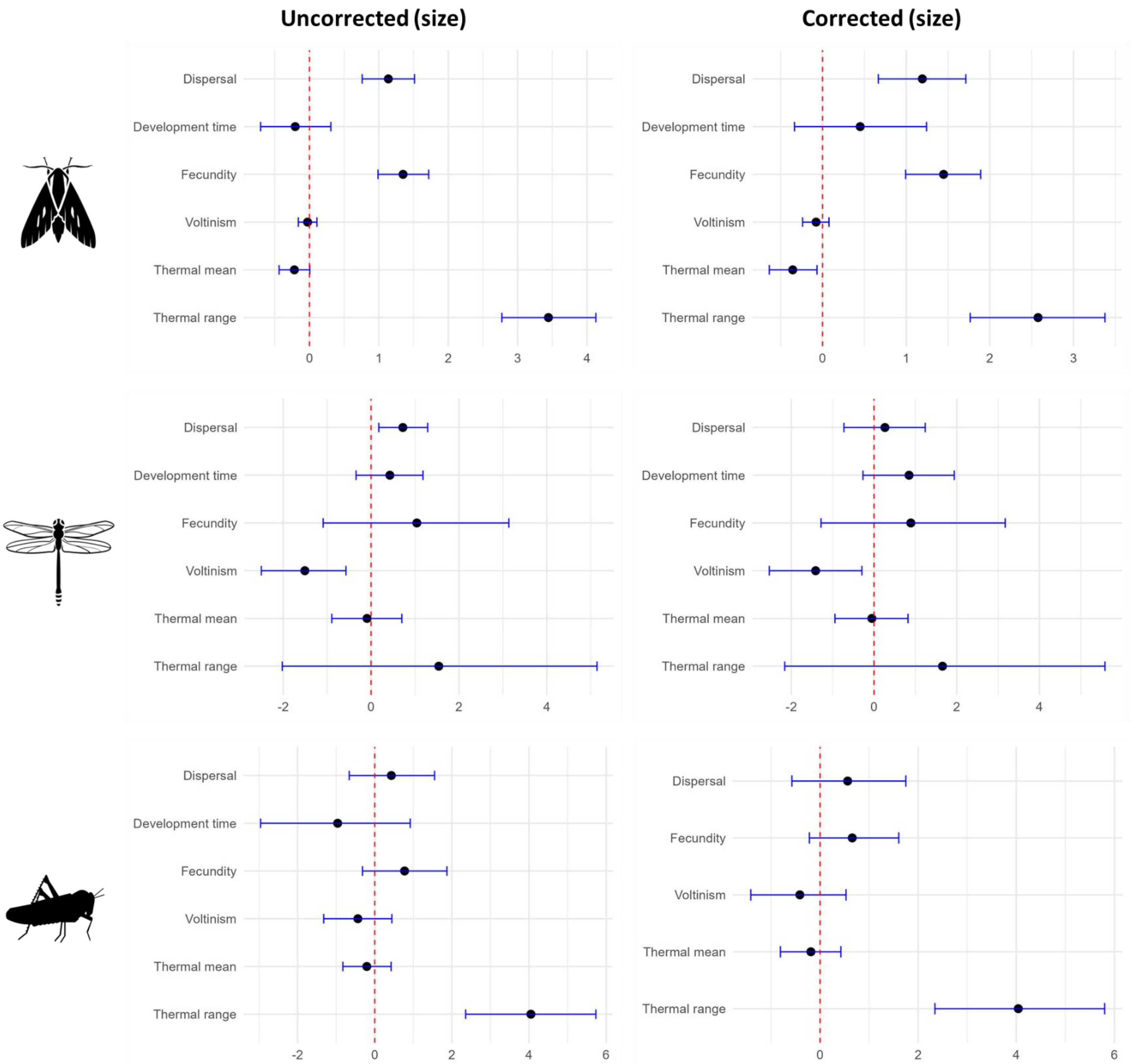

*Figure S.3.1b: comparison of uncorrected (left) and phylogenetically corrected (right) correlations with size for butterflies, dragonflies and grasshoppers. Each point represents the correlation estimate with size as the independent variable and the corresponding trait as dependent variable. The error bars represent the 95% credibility intervals.*

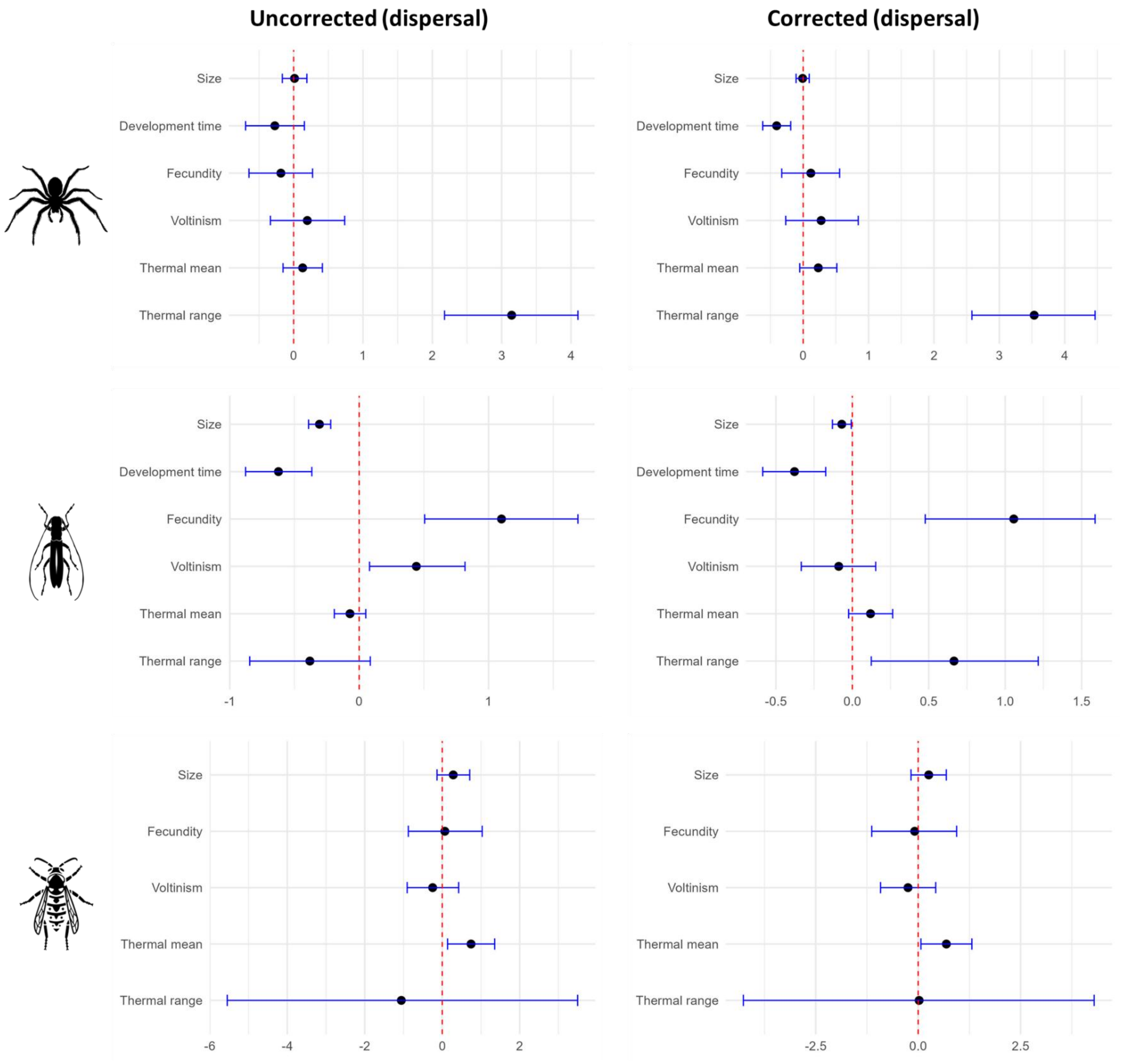

*Figure S.3.2a: comparison of uncorrected (left) and phylogenetically corrected (right) correlations with dispersal for spiders, beetles and hymenopterans. Each point represents the correlation estimate with dispersal as the independent variable and the corresponding trait as dependent variable. The error bars represent the 95% credibility intervals.*

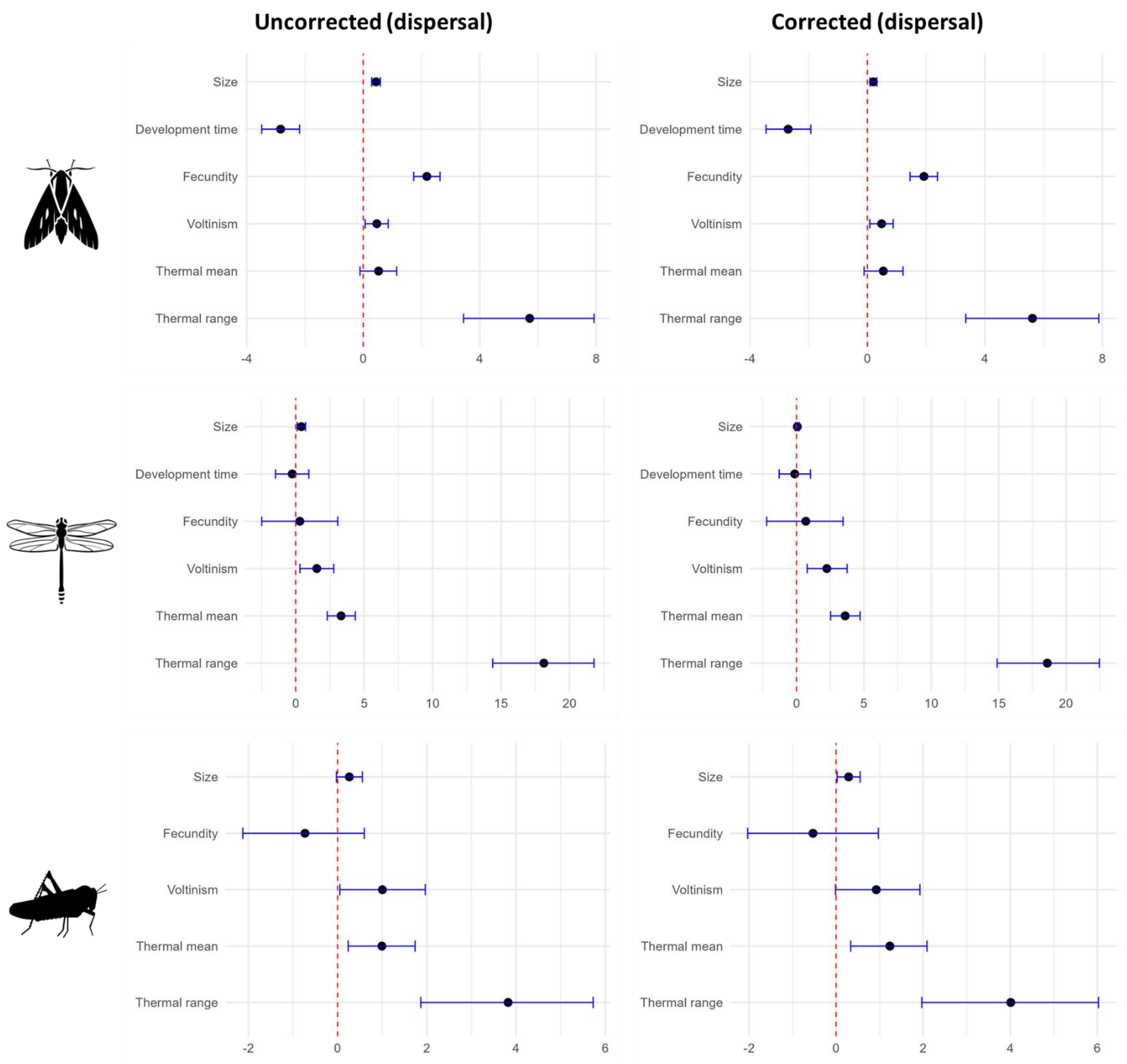

*Figure S.3.2b: comparison of uncorrected (left) and phylogenetically corrected (right) correlations with dispersal for butterflies, dragonflies and grasshoppers. Each point represents the correlation estimate with dispersal as the independent variable and the corresponding trait as dependent variable. The error bars represent the 95% credibility intervals.*

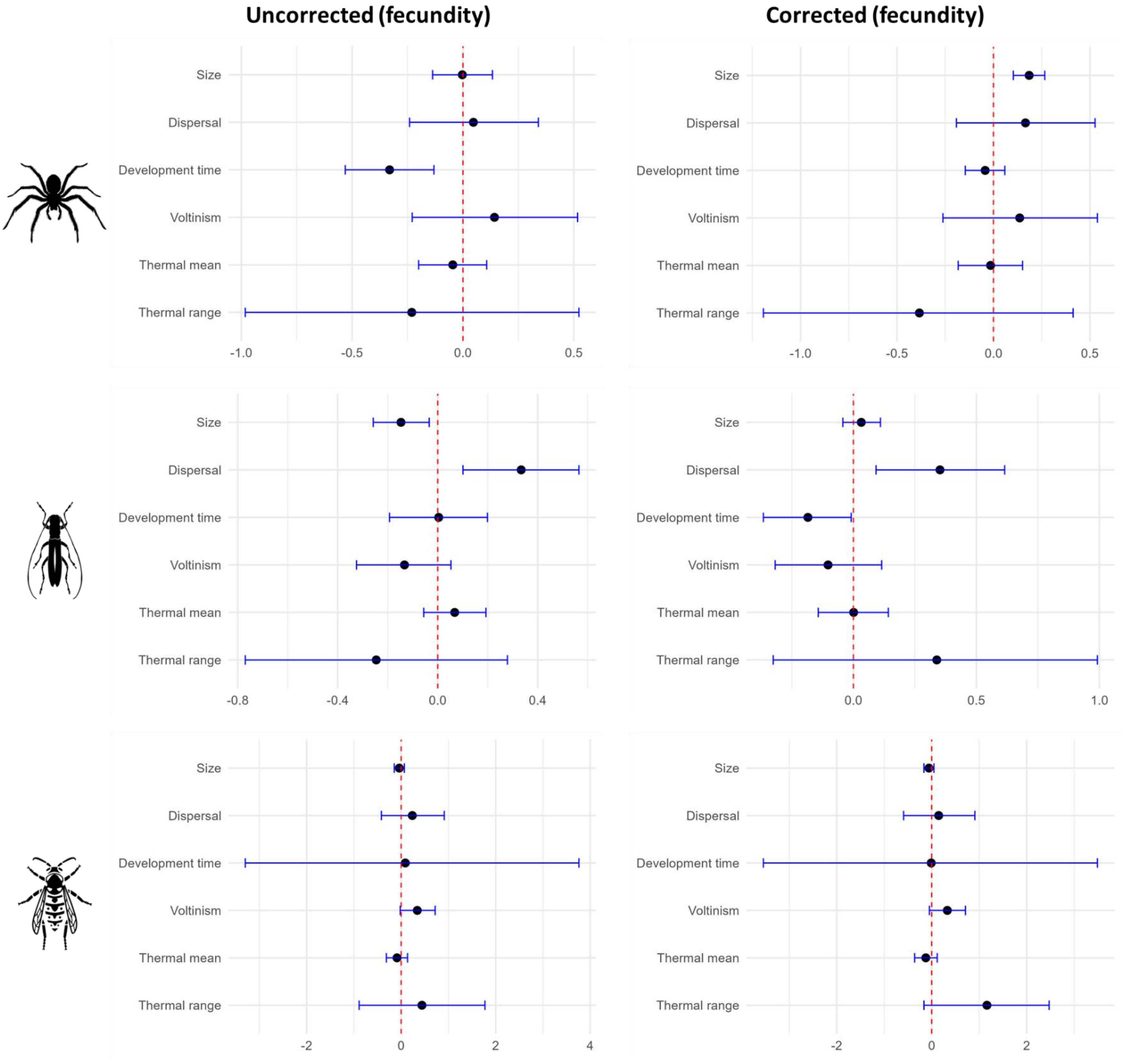

*Figure S.3.3a: comparison of uncorrected (left) and phylogenetically corrected (right) correlations with fecundity for spiders, beetles and hymenopterans. Each point represents the correlation estimate with fecundity as the independent variable and the corresponding trait as dependent variable. The error bars represent the 95% credibility intervals.*

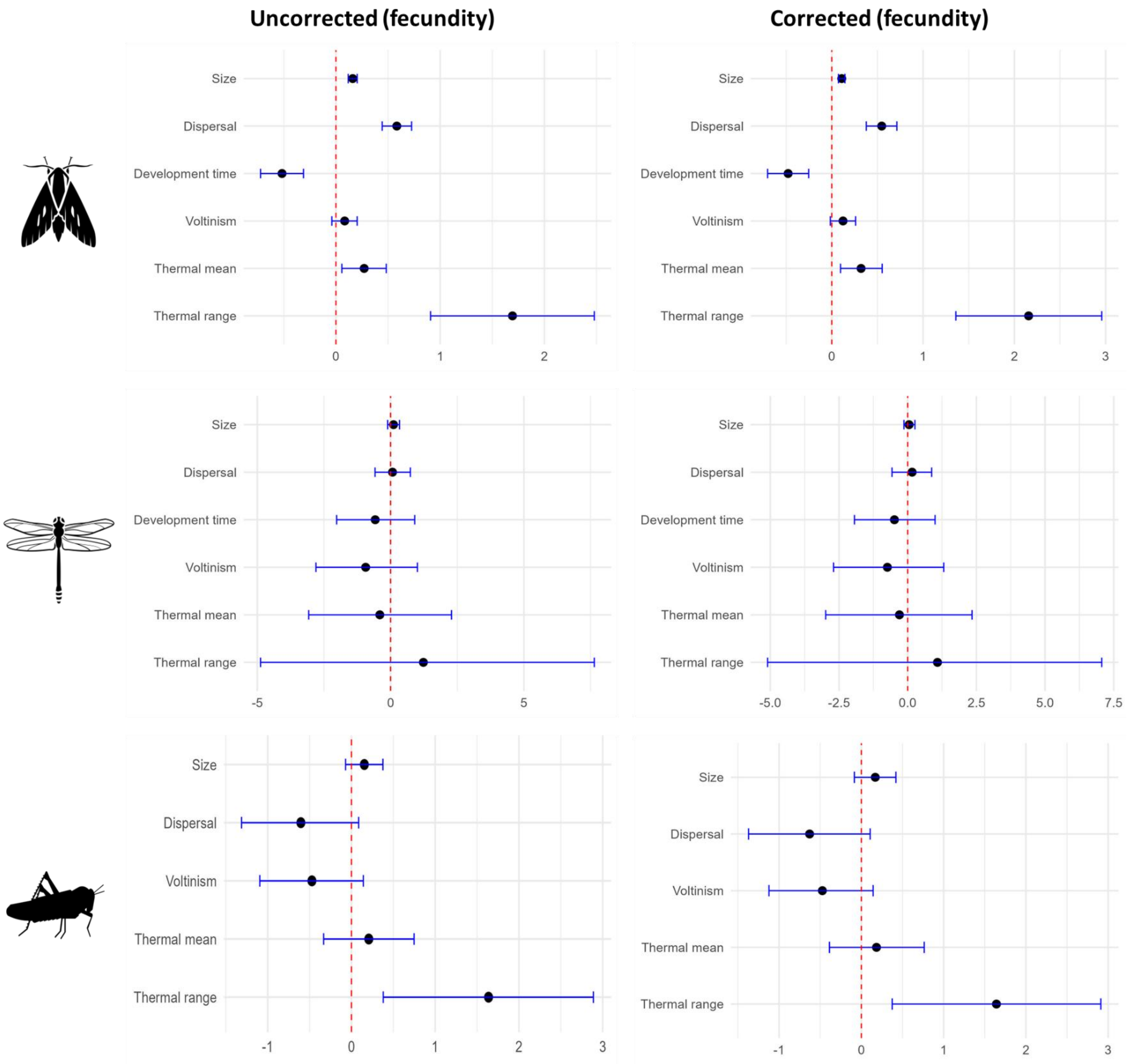

*Figure S.3.3b: comparison of uncorrected (left) and phylogenetically corrected (right) correlations with fecundity for butterflies, dragonflies and grasshoppers. Each point represents the correlation estimate with fecundity as the independent variable and the corresponding trait as dependent variable. The error bars represent the 95% credibility intervals.*

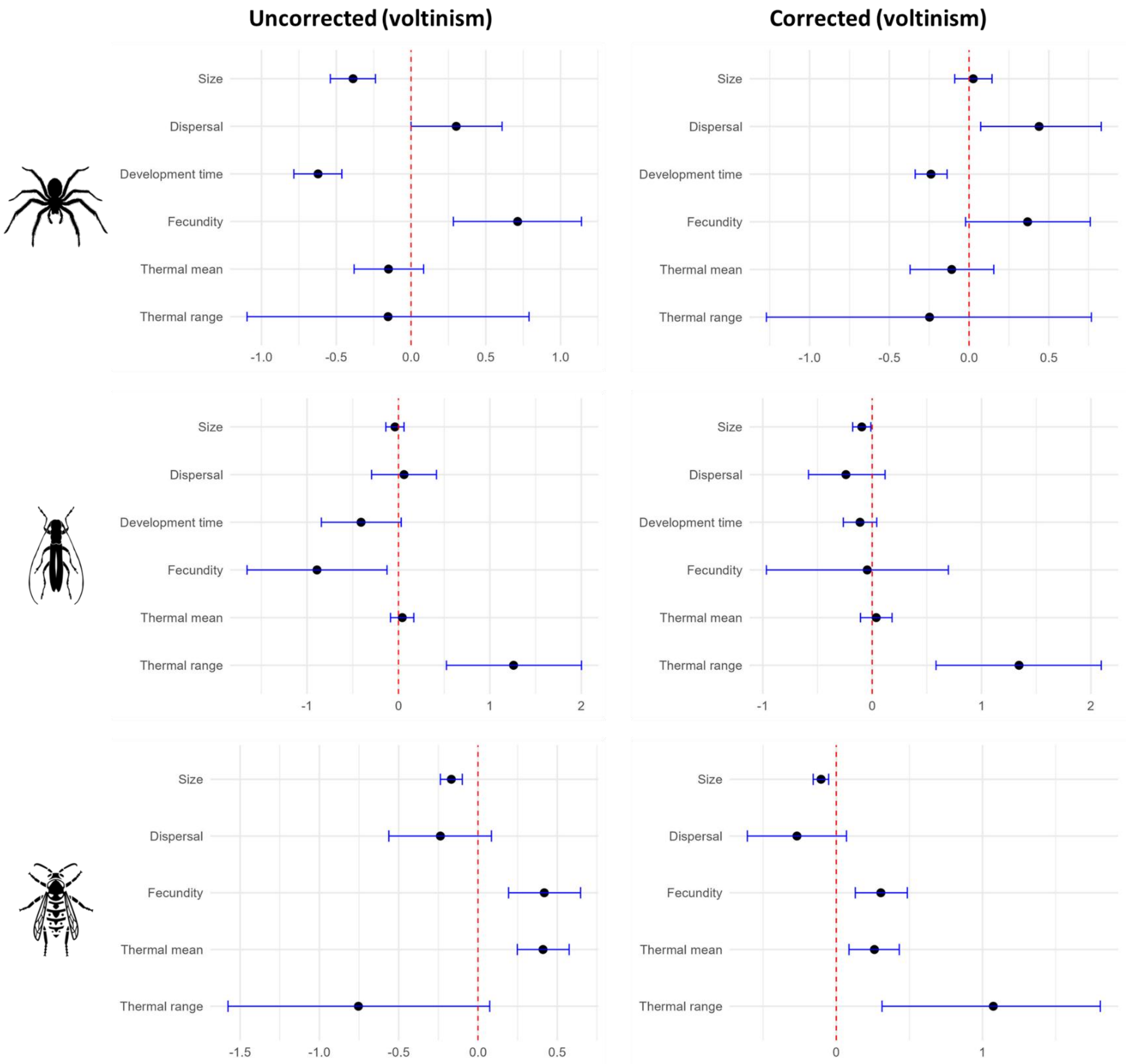

*Figure S.3.4a: comparison of uncorrected (left) and phylogenetically corrected (right) correlations with voltinism for spiders, beetles and hymenopterans. Each point represents the correlation estimate with voltinism as the independent variable and the corresponding trait as dependent variable. The error bars represent the 95% credibility intervals.*

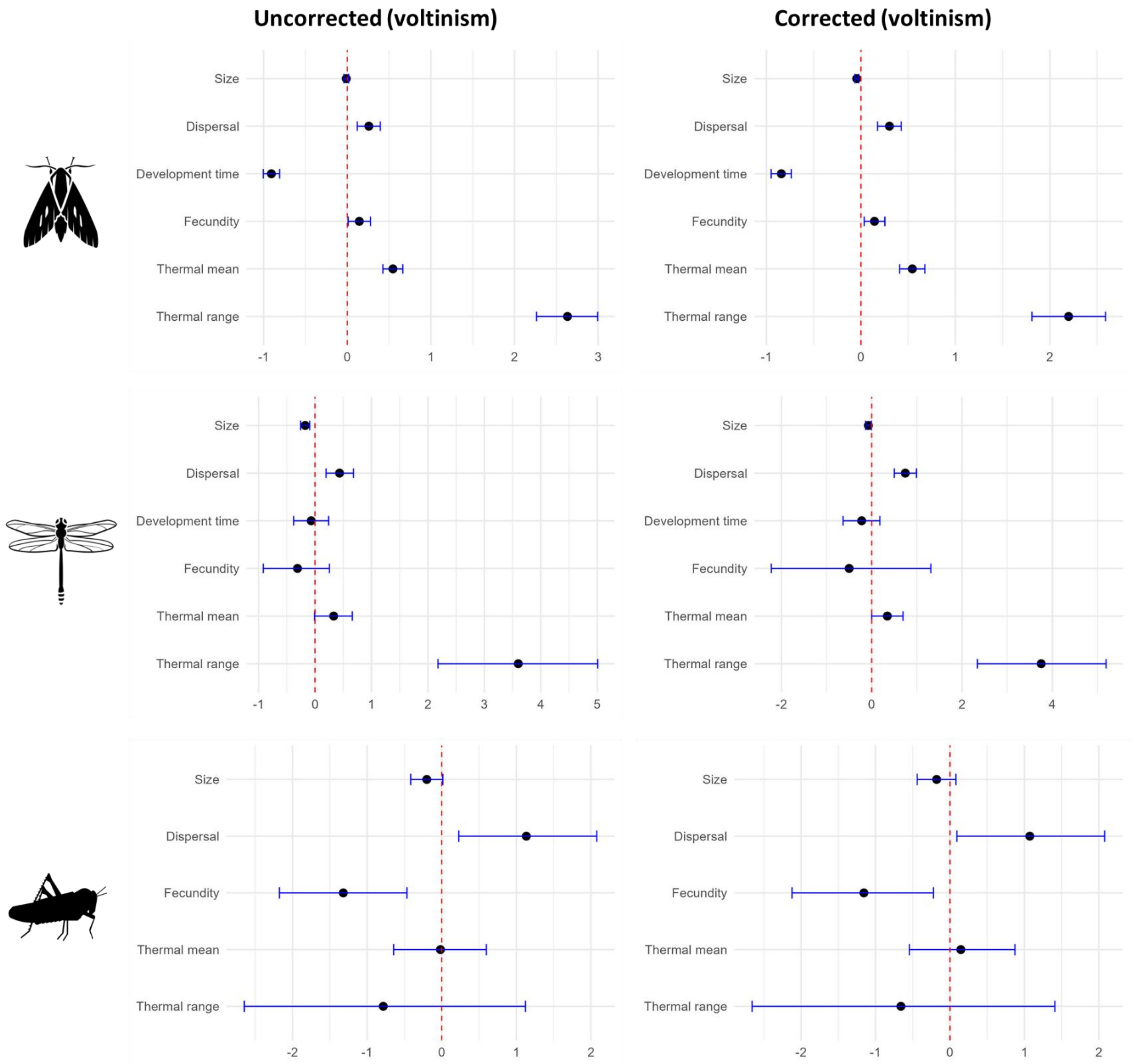

*Figure S.3.4b: comparison of uncorrected (left) and phylogenetically corrected (right) correlations with voltinism for butterflies, dragonflies and grasshoppers. Each point represents the correlation estimate with voltinism as the independent variable and the corresponding trait as dependent variable. The error bars represent the 95% credibility intervals.*

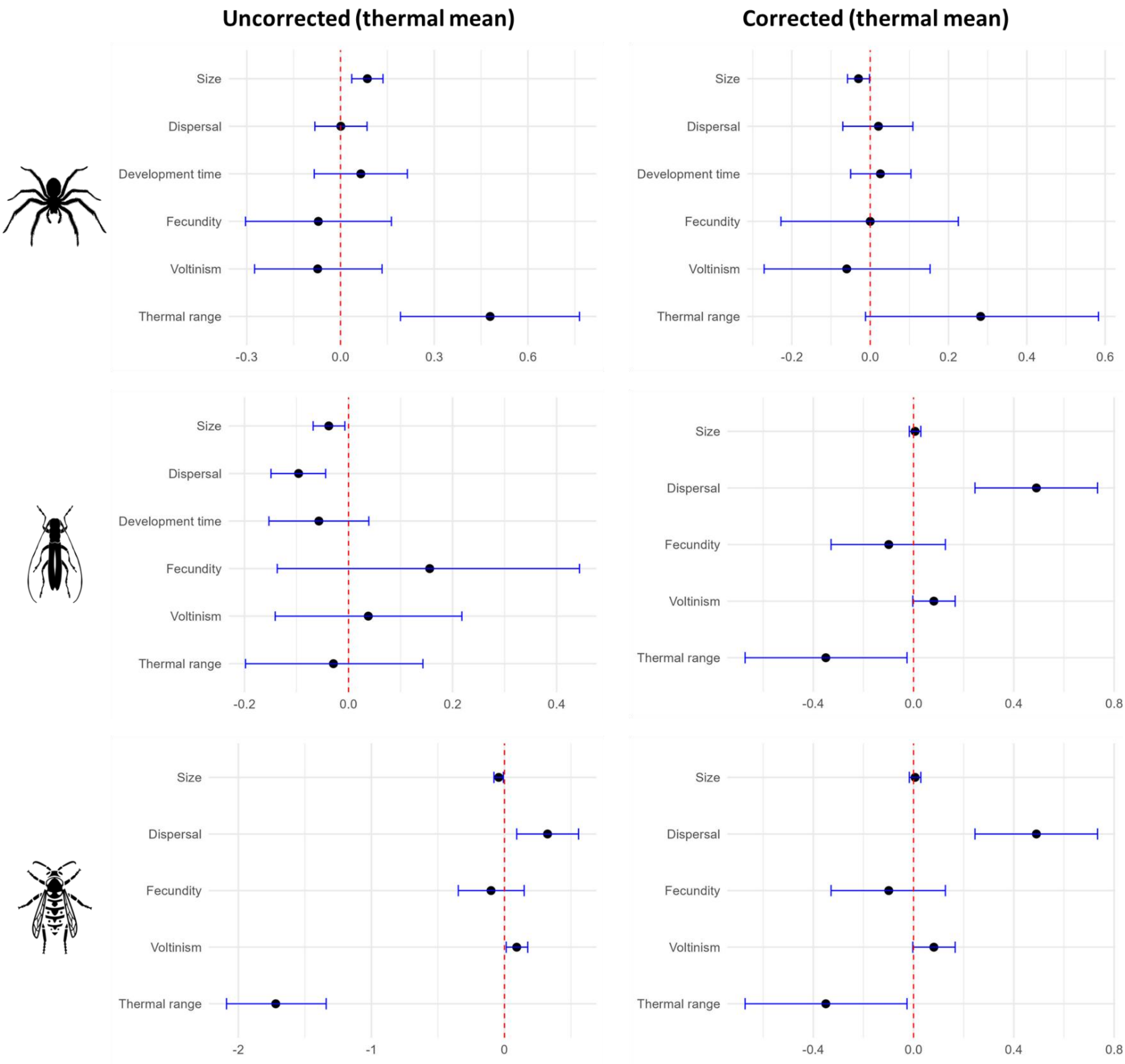

*Figure S.3.5a: comparison of uncorrected (left) and phylogenetically corrected (right) correlations with thermal mean for spiders, beetles and hymenopterans. Each point represents the correlation estimate with thermal mean as the independent variable and the corresponding trait as dependent variable. The error bars represent the 95% credibility intervals.*

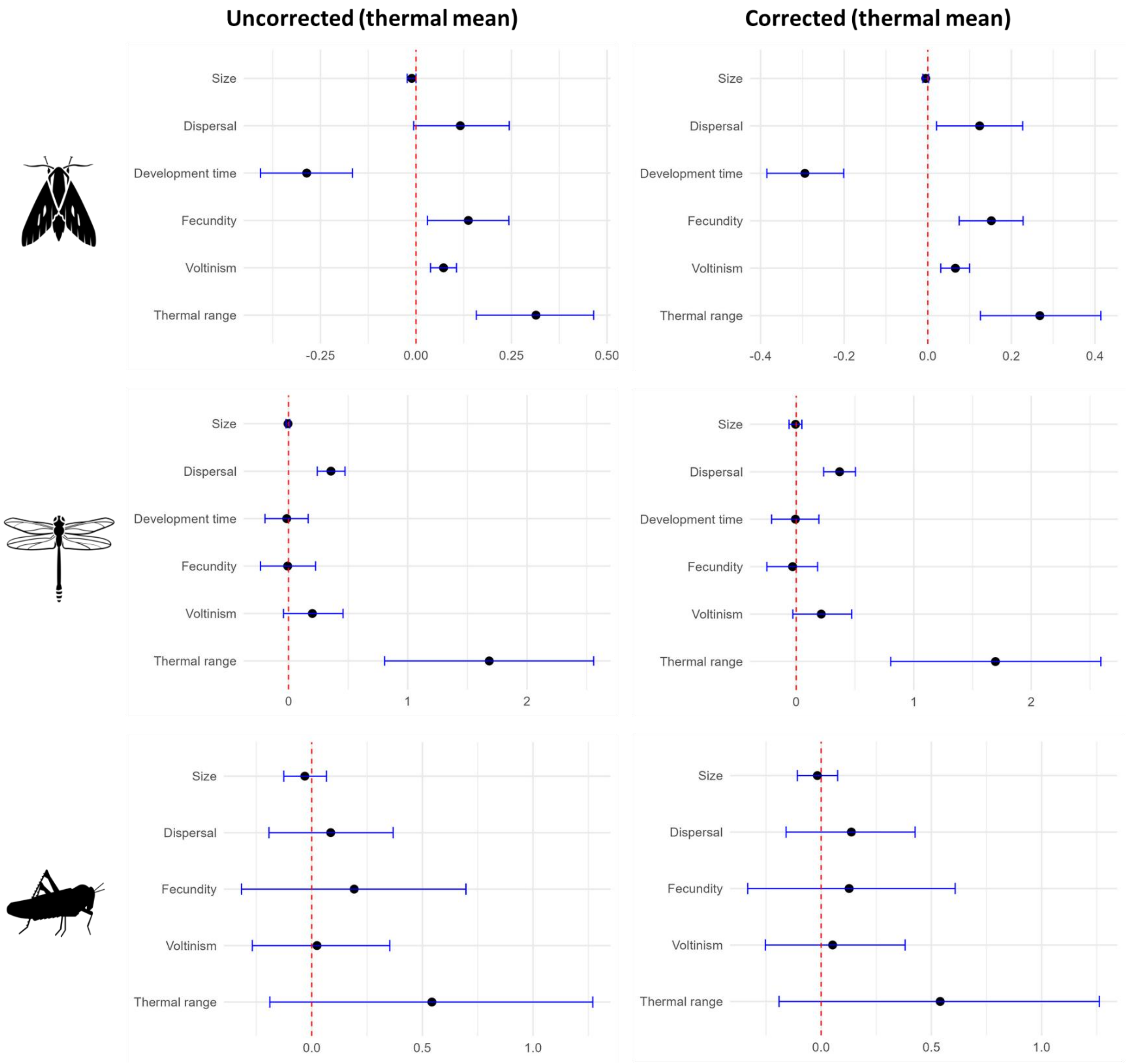

*Figure S.3.5b: comparison of uncorrected (left) and phylogenetically corrected (right) correlations with thermal mean for butterflies, dragonflies and grasshoppers. Each point represents the correlation estimate with thermal mean as the independent variable and the corresponding trait as dependent variable. The error bars represent the 95% credibility intervals.*

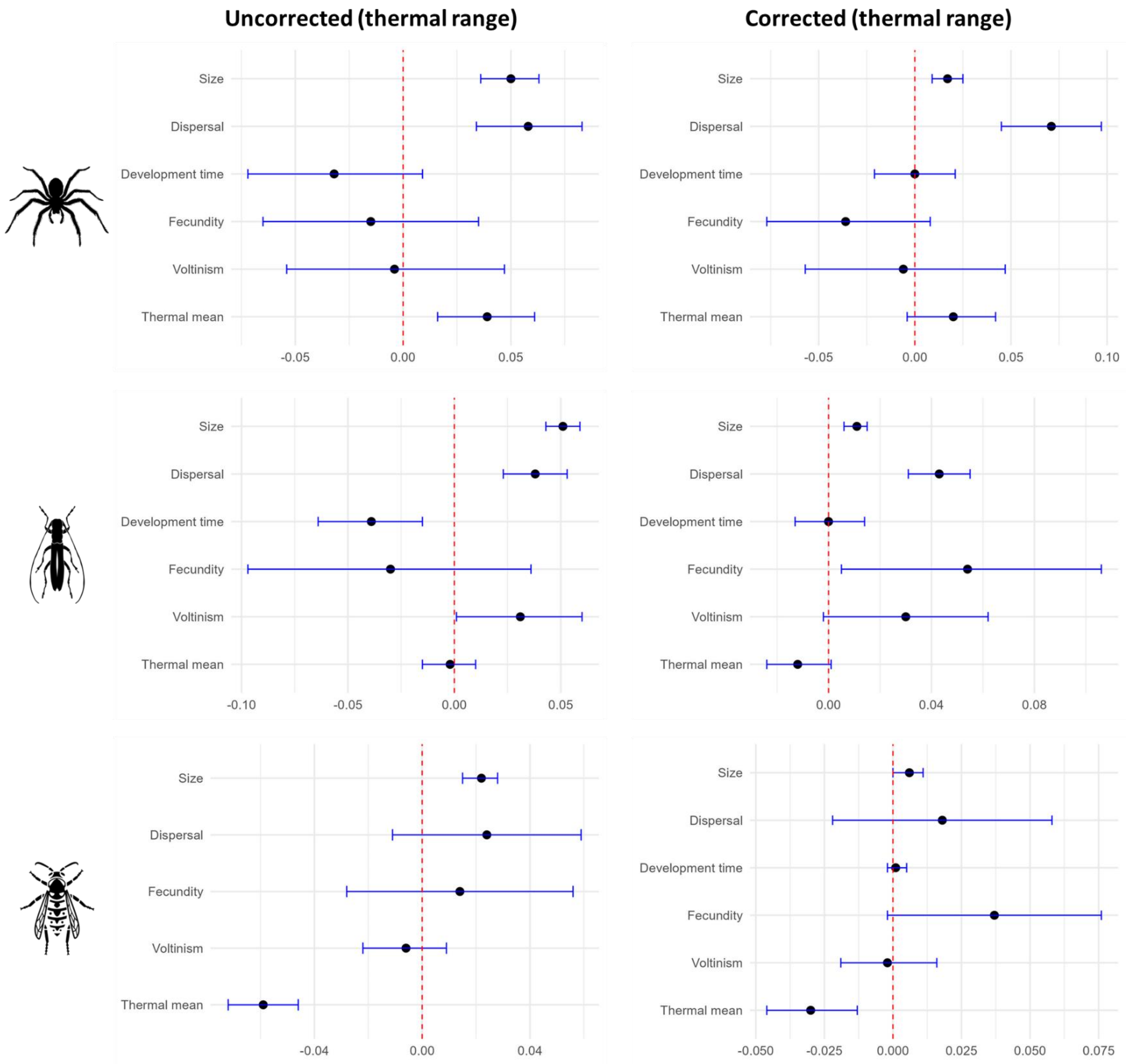

*Figure S.3.6a: comparison of uncorrected (left) and phylogenetically corrected (right) correlations with thermal range for spiders, beetles and hymenopterans. Each point represents the correlation estimate with thermal range as the independent variable and the corresponding trait as dependent variable. The error bars represent the 95% credibility intervals.*

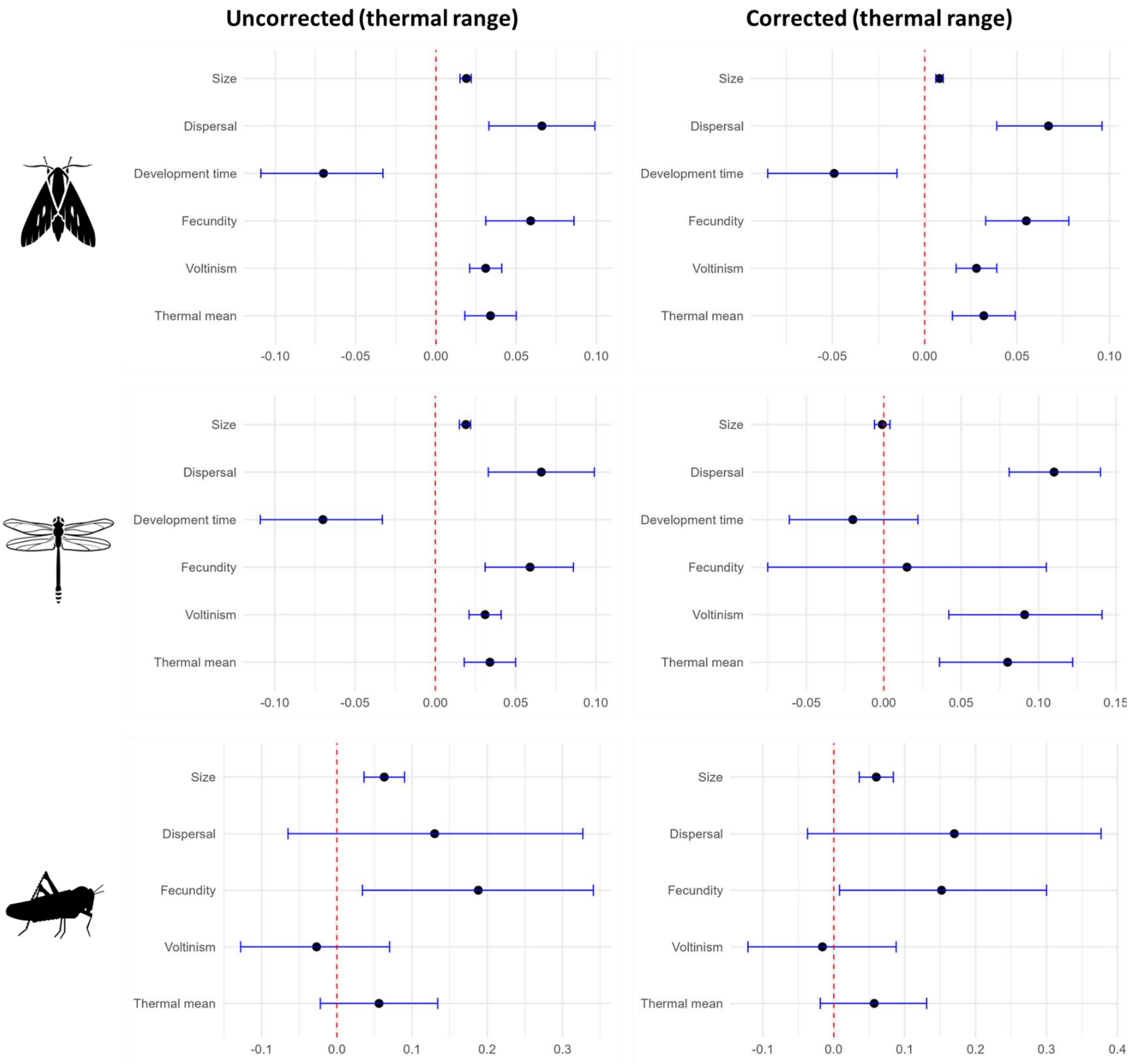

*Figure S.3.6b: comparison of uncorrected (left) and phylogenetically corrected (right) correlations with thermal range for butterflies, dragonflies and grasshoppers. Each point represents the correlation estimate with thermal range as the independent variable and the corresponding trait as dependent variable. The error bars represent the 95% credibility intervals.*

#### S.4 Correlation matrices for the different taxonomic groups

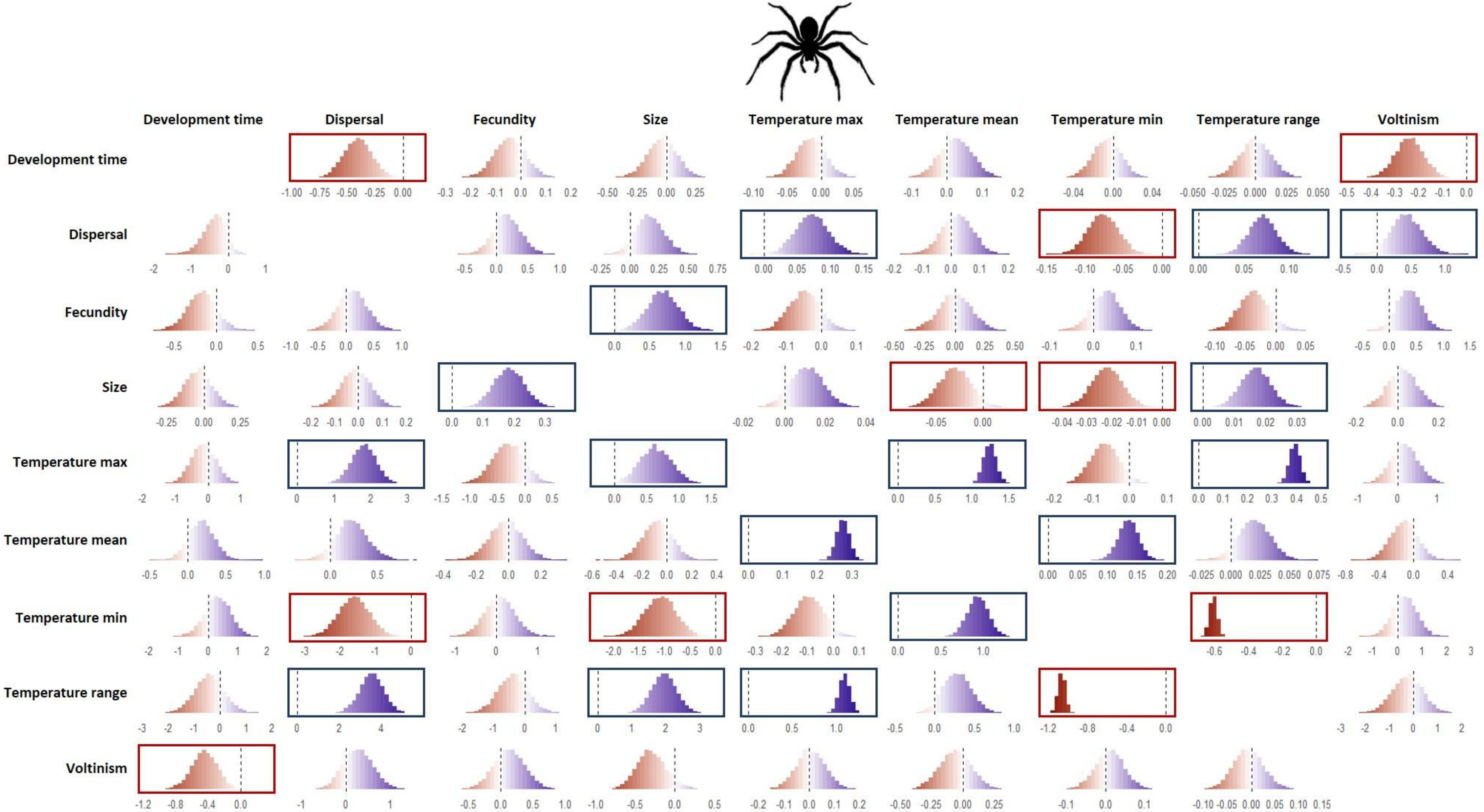

*Figure S.4.1: Pairwise posterior distributions for trait covariations (slopes of the linear regressions) within Araneae. Column names represent independent variables, while row names denote dependent variables Negative covariations are highlighted in red and positive covariations in blue. Effect sizes that are clearly determined (i.e. the 95% interval does not include zero) are outlined in the corresponding colour. Icon at the top: Spider by Matthew Davis from Noun Project (CC BY-NC-ND 2.0).*

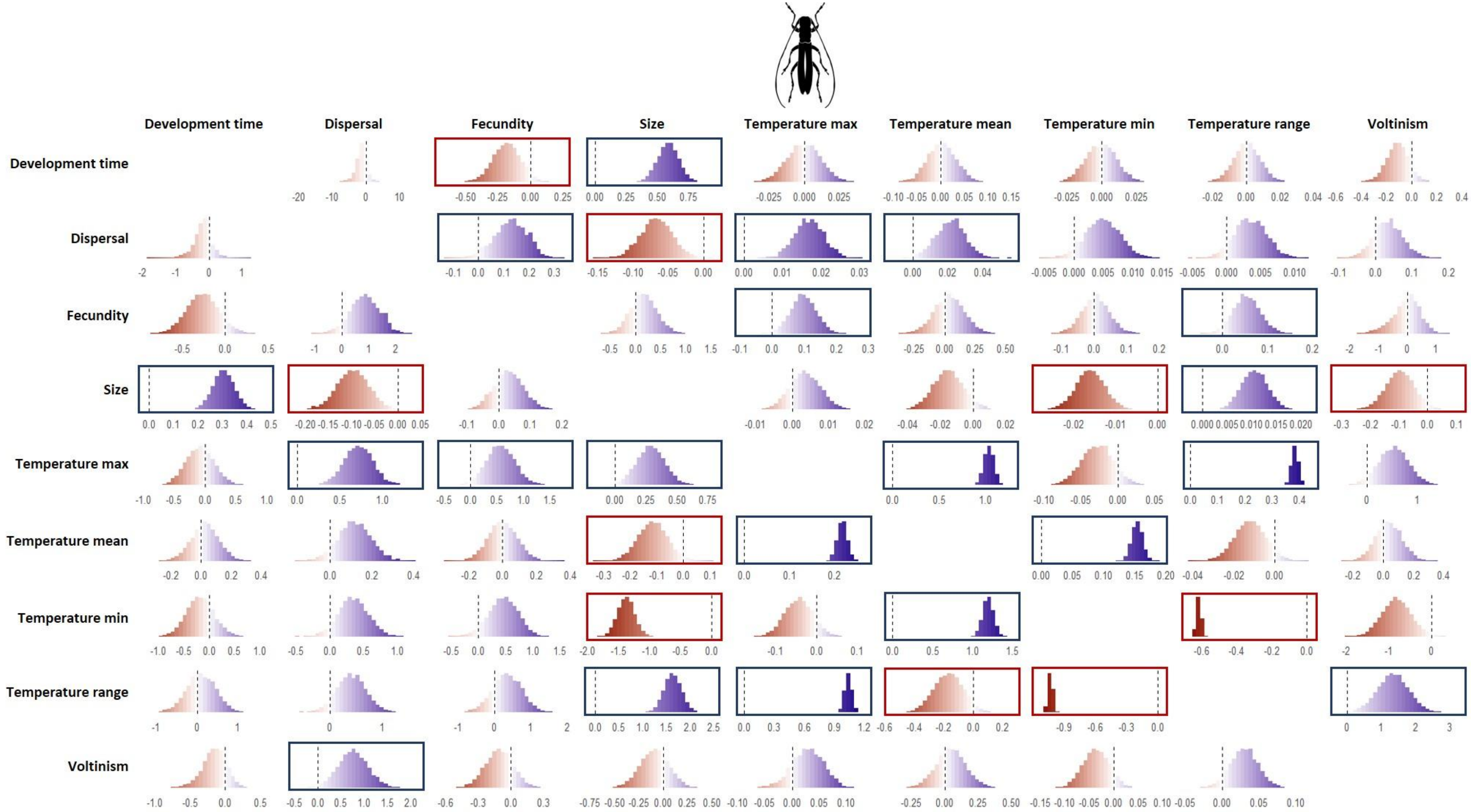

*Figure S.4.2: Pairwise posterior distributions for trait covariations (slopes of the linear regressions) within Coleoptera. Column names represent independent variables, while row names denote dependent variables Negative covariations are highlighted in red and positive covariations in blue. Effect sizes that are clearly determined (i.e. the 95% interval does not include zero) are outlined in the corresponding colour. Icon at the top: Beetle by Rachel Siao from Noun Project (CC BY-NC-ND 2.0).*

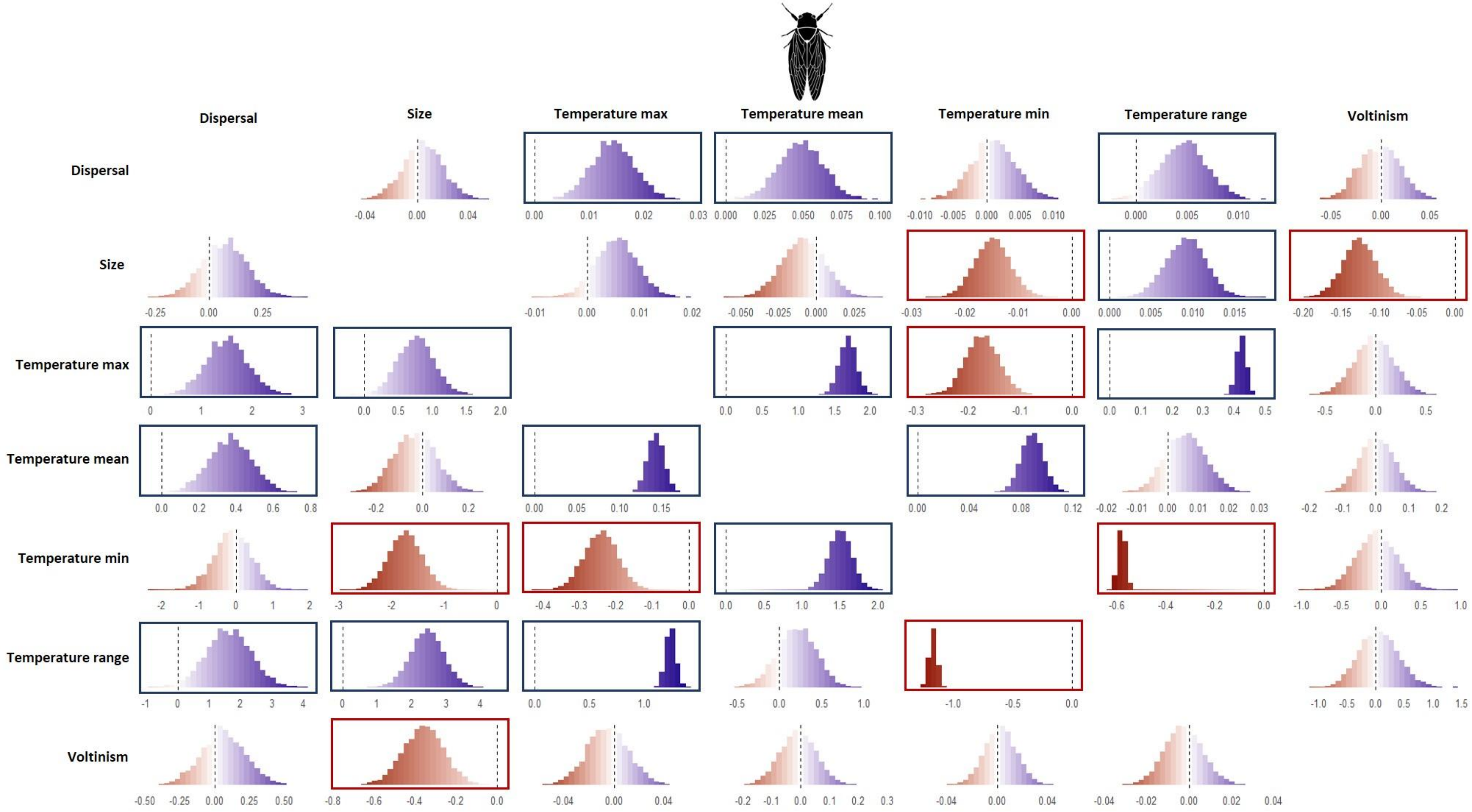

*Figure S.4.3: Pairwise posterior distributions for trait covariations (slopes of the linear regressions) within Hemiptera. Column names represent independent variables, while row names denote dependent variables Negative covariations are highlighted in red and positive covariations in blue. Effect sizes that are clearly determined (i.e. the 95% interval does not include zero) are outlined in the corresponding colour. Icon at the top: Cicada by Alejandro Capellan from Noun Project (CC BY-NC-ND 2.0).*

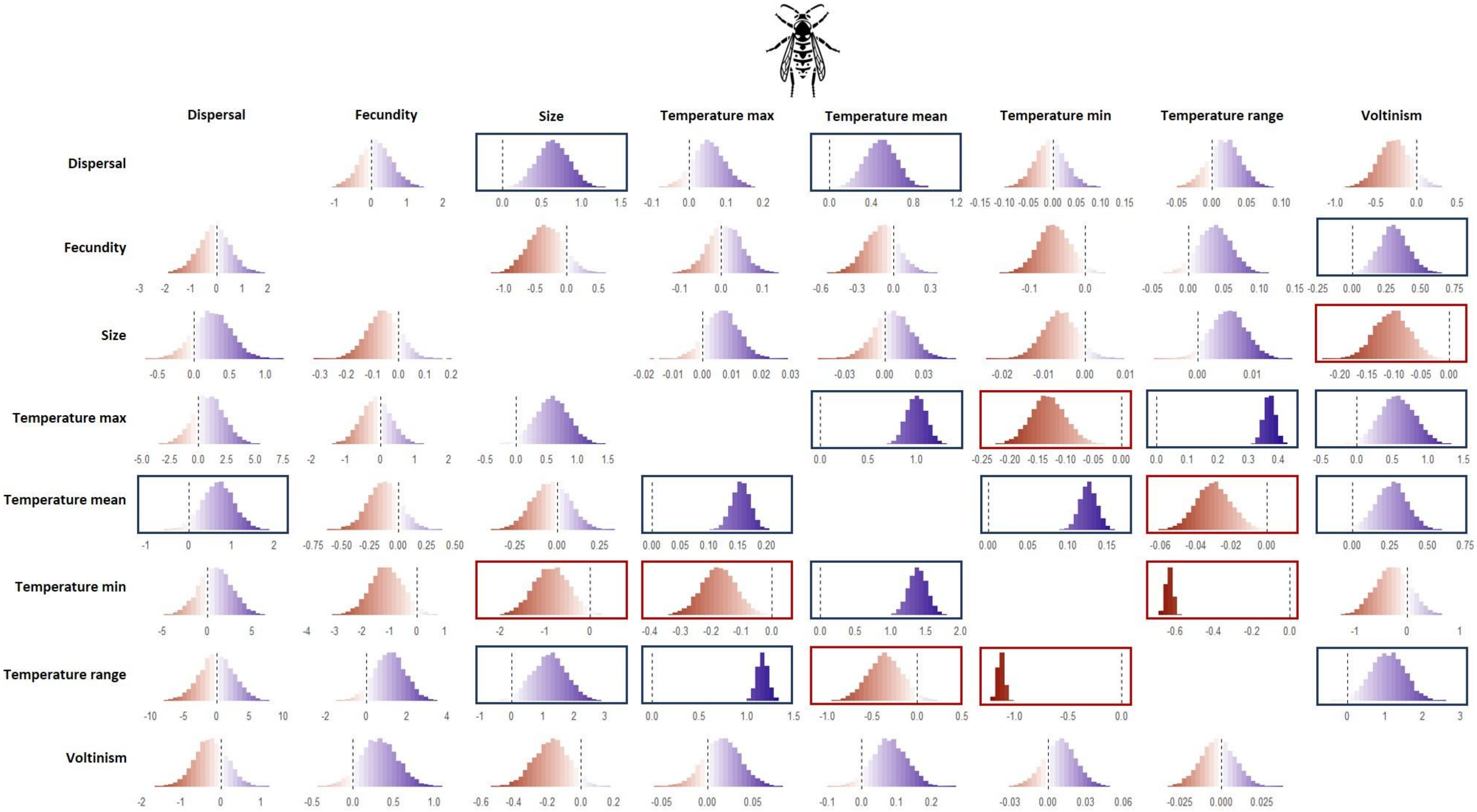

*Figure S.4.4: Pairwise posterior distributions for trait covariations (slopes of the linear regressions) within Hymenoptera. Column names represent independent variables, while row names denote dependent variables Negative covariations are highlighted in red and positive covariations in blue. Effect sizes that are clearly determined (i.e. the 95% interval does not include zero) are outlined in the corresponding colour. Icon at the top: wasp by parkijsun from Noun Project (CC BY-NC-ND 2.0).*

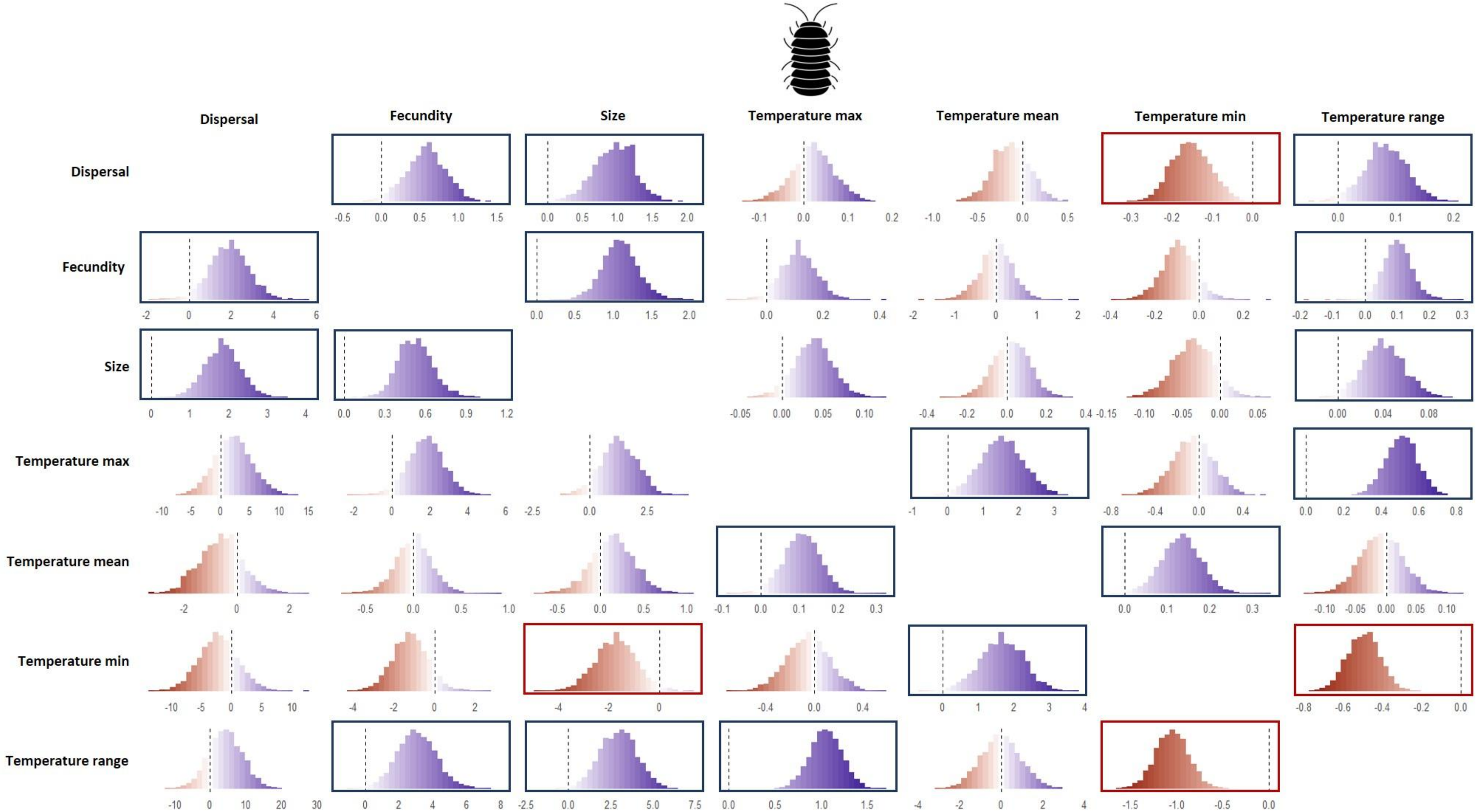

*Figure S.4.5: Pairwise posterior distributions for trait covariations (slopes of the linear regressions) within Isopoda. Column names represent independent variables, while row names denote dependent variables Negative covariations are highlighted in red and positive covariations in blue. Effect sizes that are clearly determined (i.e. the 95% interval does not include zero) are outlined in the corresponding colour. Icon at the top: isopod by Pham Thanh Lôc from Noun Project (CC BY-NC-ND 2.0).*

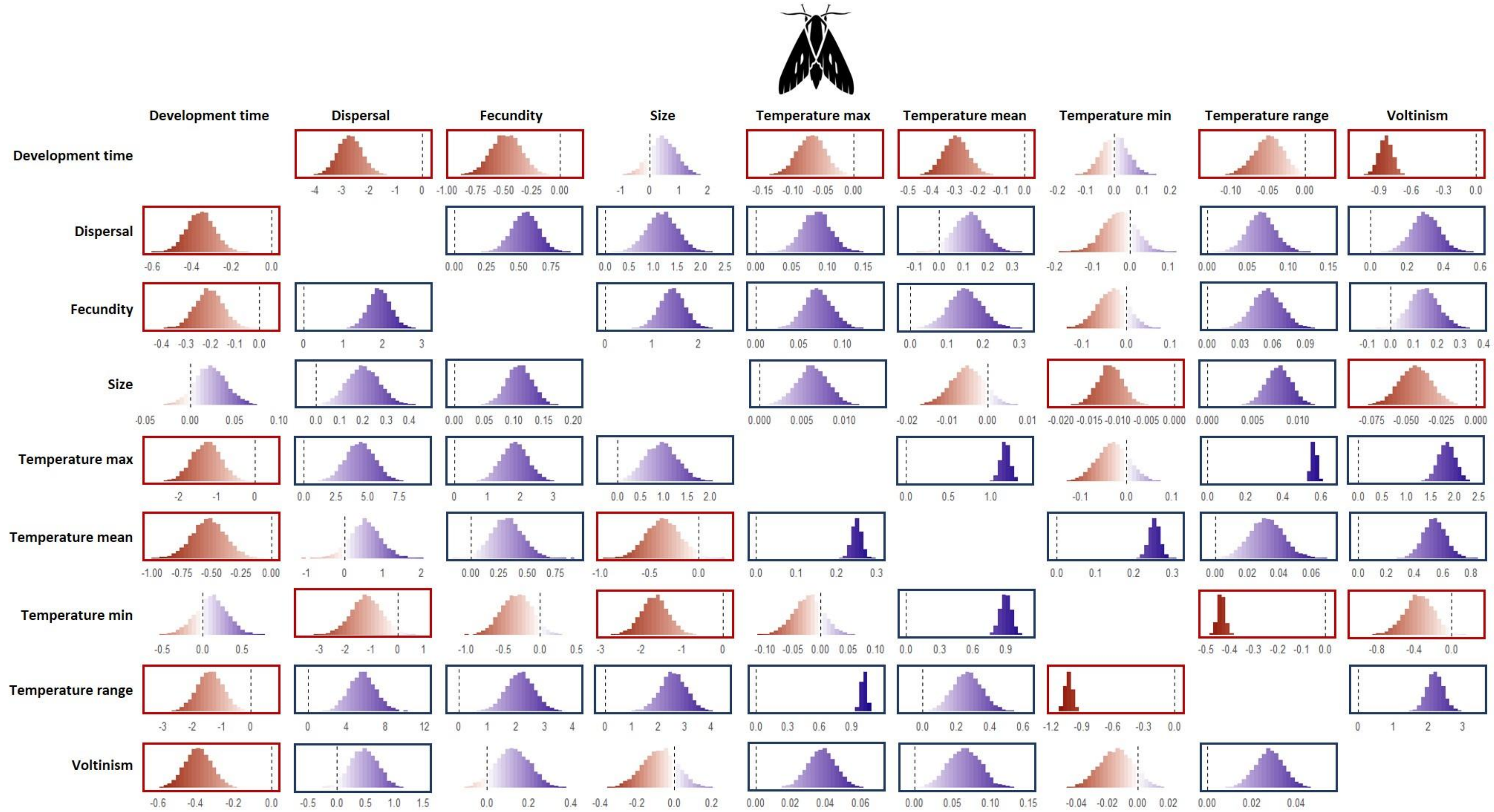

*Figure S.4.6: Pairwise posterior distributions for trait covariations (slopes of the linear regressions) within Lepidoptera. Column names represent independent variables, while row names denote dependent variables Negative covariations are highlighted in red and positive covariations in blue. Effect sizes that are clearly determined (i.e. the 95% interval does not include zero) are outlined in the corresponding colour. Icon at the top: Moth by parkijsun from Noun Project (CC BY-NC-ND 2.0).*

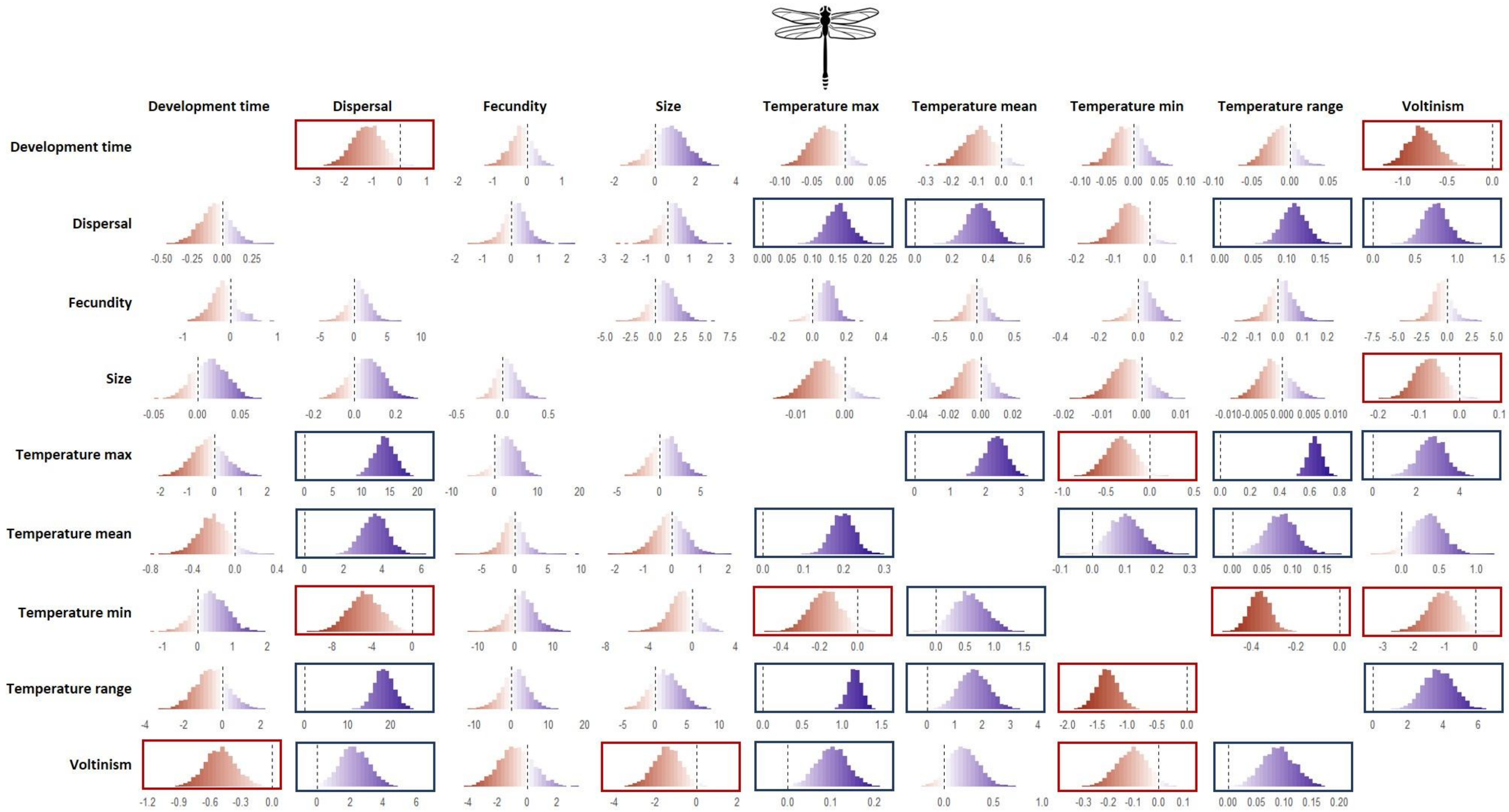

*Figure S.4.7: Pairwise posterior distributions for trait covariations (slopes of the linear regressions) within Odonata. Column names represent independent variables, while row names denote dependent variables Negative covariations are highlighted in red and positive covariations in blue. Effect sizes that are clearly determined (i.e. the 95% interval does not include zero) are outlined in the corresponding colour. Icon at the top: Dragonfly nu Hermine Blanquart from Noun Project (CC BY-NC-ND 2.0).*

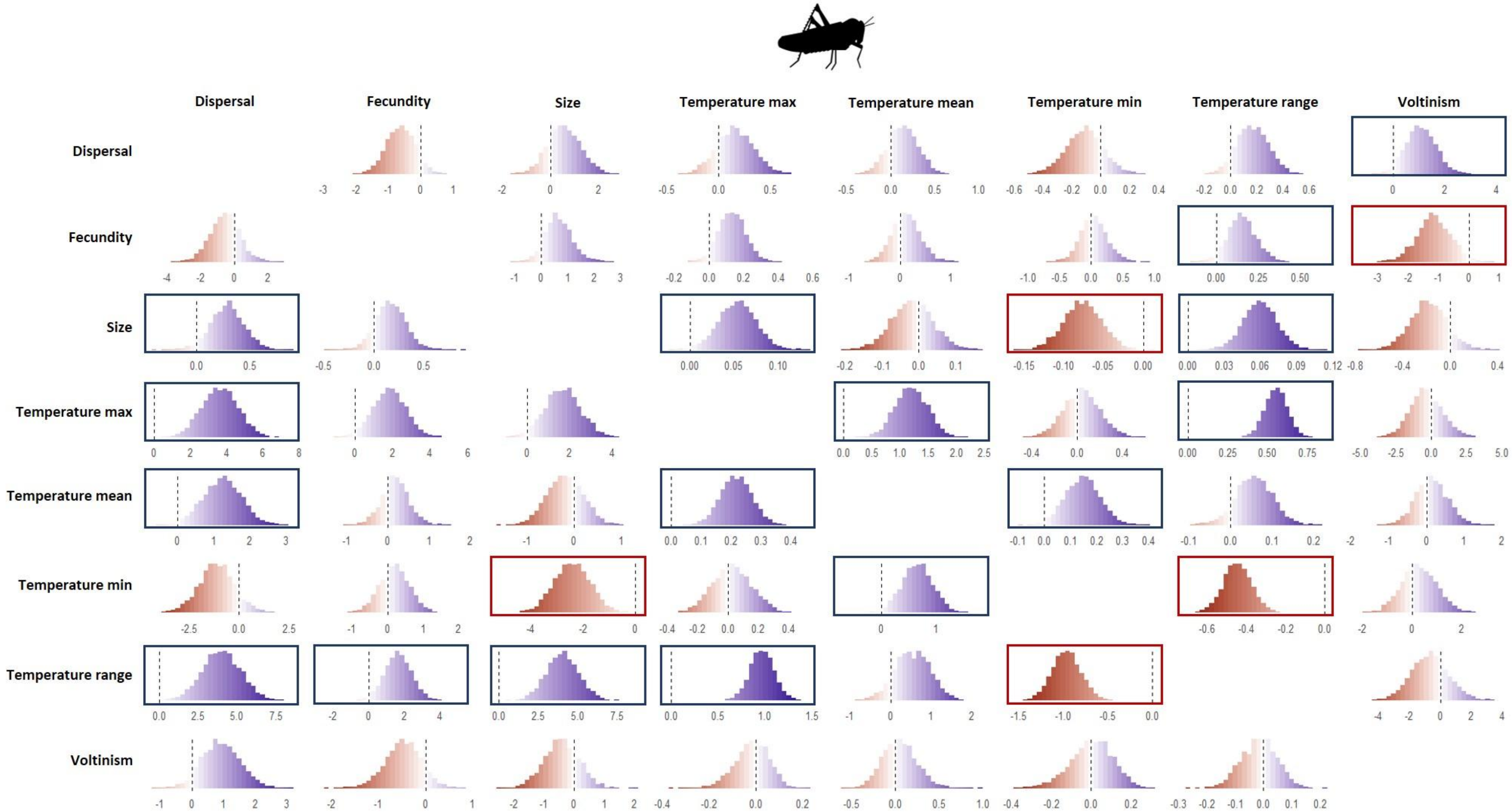

*Figure S.4.8: Pairwise posterior distributions for trait covariations (slopes of the linear regressions) within Orthoptera. Column names represent independent variables, while row names denote dependent variables Negative covariations are highlighted in red and positive covariations in blue. Effect sizes that are clearly determined (i.e. the 95% interval does not include zero) are outlined in the corresponding colour. Icon at the top: Cricket by Ed Harrison from Noun Project (CC BY-NC-ND 2.0).*

#### S.5 Phylogenetic Principal Component Analysis

*Table S.5: Output of the phylogenetic PCA. Loadings represent the correlation coefficients between each variable and the principal components. Standard deviation denotes the square root of each principal component's eigenvalue, reflecting the amount of variance each component captures. Proportion of variance specifies the percentage of total variance explained by each component, while cumulative proportion of variance indicates the cumulative variance accounted for by the addition of each subsequent component.*

| | PC1 | PC2 | PC3 | PC4 |
|---|---|---|---|---|
| **Variable loadings** | | | | |
| Size | -0.332 | 0.805 | -0.002 | 0.252 |
| Fecundity | 0.293 | 0.639 | 0.053 | -0.582 |
| Development time | -0.817 | 0.297 | 0.115 | 0.127 |
| Dispersal | 0.614 | 0.170 | 0.419 | -0.220 |
| Voltinism | 0.778 | 0.008 | -0.187 | 0.235 |
| Thermal mean | 0.424 | 0.345 | -0.695 | 0.248 |
| Thermal range | 0.405 | 0.161 | 0.644 | 0.506 |
| Standard deviation | 1.480 | 1.148 | 1.060 | 0.916 |
| Proportion of variance | 0.313 | 0.188 | 0.161 | 0.120 |
| Cumulative proportion of variance | 0.313 | 0.501 | 0.662 | 0.782 |
